# Unveiling the intraspecific genomic diversity of South American lowland maize landraces

**DOI:** 10.64898/2026.10.17.759863

**Authors:** Renata F. Carvalho, Natália Carolina de Almeida Silva, Rafael Vidal, Magdalena Vaio

**Affiliations:** Facultad de Agronomía, Universidad de la República (UdelaR), Montevideo, 12900 Uruguay; Instituto Tecnológico Regional Centro Sur, Universidad Tecnológica del Uruguay (UTEC), Durazno 97000, Uruguay

**Keywords:** Genome size, heterochromatic knobs, races, *Zea mays* subsp. *mays*, in situ hybridization, repetitive sequences

## Abstract

**Background and Aims:** South American Lowland maize landraces constitute an important reservoir of the species diversity, although research regarding the territories of Uruguay and Brazil remains comparatively unexplored. This study aimed to evaluate the intragenomic variability of maize landraces from Uruguay and Brazil.

**Methods:** We estimated the genome size, number of heterochromatic knobs, and presence of B chromosomes in 15 maize landraces representing 13 races, and finally, we conducted an analysis of repetitive sequences in landraces from both countries.

**Key Results:** Regarding DNA content, a north-to-south gradient of decreasing genome size was found, significantly associated with latitude. At the knob level, a model incorporating the Country × Race interaction detected significant effects, revealing a reversal of the pattern of knob number per race among countries. B chromosomes were found only in Uruguayan races, whereas regarding repetitive sequences, a trend of increasing retrotransposons and decreasing knobs was observed as DNA content decreased.

**Conclusions:** Taken together, these results suggest that genome size may be influenced by selective pressures associated with local adaptation, although further investigation is needed to test this hypothesis. Furthermore, the knob profiles of specific races across countries, as well as the presence of B chromosomes in Uruguay, may reflect distinct histories of dispersal, local selection, and even introgression, highlighting the complexity of the processes shaping maize genomic architecture across geographic gradients.

## Introduction

Maize (*Zea mays* subsp. *mays* L.) is an important staple crop worldwide, and two of the world’s largest maize producers are located in the Americas: the United States and Brazil (FAO 2026). This crop has been classified into races and landraces due to its extensive genetic diversity. A maize race was originally defined as a group of individuals sharing certain characteristics that allow them to be recognized as a distinct group (Anderson and Cutler 1942). Landraces are genetically dynamic populations characterized by high variability resulting from selection, conservation, and management practices carried out by traditional farmers and Indigenous peoples (Silva *et al*. 2020, 2021). Therefore, a maize race can comprise multiple landraces. Approximately 300 maize races have been described in the literature, of which 260 occur in Latin America, most of them in South America (Salhuana and Pollak 2006).

Two genetic groups with distinct geographic distributions have been identified among extant maize landraces in South America: the Andean and Tropical Lowland groups (Vigouroux *et al*. 2008). A recent review of the history of maize in these regions highlighted the importance of cytogenetic studies and research on adaptation, diversity, and genetic structure for understanding South American maize. However, the authors emphasized the limited information available on the adaptive processes experienced by lowland landraces (Dominguez *et al*. 2024).

In the South American lowlands, which include areas below 1,500 m above sea level, studies on maize races and landraces had not been updated since 1977 (Paterniani and Goodman 1977). This gap prompted a collective effort to map, identify, and classify the diversity of maize landraces conserved *in situ* on-farm in Uruguay and Brazil. These studies provided a new perspective on maize diversity in these countries, identifying a total of 29 races and three race complexes based on ethnobotanical, phenotypic, and molecular characteristics (Silva *et al*. 2020, 2021; Costa *et al*. 2022). Based on the resulting database, subsequent studies have provided a deeper understanding of the conservation status of maize (Costa *et al*. 2022), its dispersal through human migrations and subsequent diversification into different races and landraces (Kistler *et al*. 2020; Costa *et al*. 2022), the identification of microcenters and regions of genetic diversity (Costa *et al*. 2023), and the identification of archaeological samples with primitive maize characteristics far from its center of origin (Costa *et al*. 2024). These findings reinforce the relevance of lowland germplasm to the history of maize, demonstrating its dynamic nature and ongoing evolutionary processes, which challenge the hypothesis of genetic erosion through replacement by commercial cultivars (Gimenes and Lopes 2000; Silva *et al*. 2020).

Historically, cytogenetic characterization has been an important component of maize race identification, including the determination of the number and location of chromosomal knobs, the presence of B chromosomes, and the organization and structure of the karyotype. The position and number of knobs vary widely among races and species of the genus *Zea*, and the observed patterns have been used by several authors to characterize, identify, and classify races, as well as to suggest relationships among them (Wellhausen *et al*. 1952; McClintock *et al*. 1981; Hanson 1984; Gonzalez *et al*. 2013; Kato 2023). In addition, DNA content quantification has made it possible to accurately estimate and compare genome sizes among maize landraces and races, as well as among different species of the genus *Zea* (Rayburn *et al*. 1985; Realini *et al*. 2016b). For instance, Laurie and Bennett (1985) reported that *Z. perennis* (2n = 40) exhibited the highest DNA content (2C = 10.56 pg), followed by *Z. luxurians* (2C = 9.14 pg) and *Z. diploperennis* (2C = 5.28 pg). For *Z. mays* subsp. *mays*, the genome size ranged from 2C = 4.92 pg to 2C = 6.74 pg. Intraspecific variation has also been widely documented: maize races from northern Argentina exhibited values ranging from 2C = 5.00 to 2C = 6.75 pg (Poggio *et al*. 1998), whereas Mexican landraces along an altitudinal gradient showed significant variation, ranging from 2C = 5.92 pg to 2C = 7.10 pg (Díez *et al*. 2013). Similarly, significant variation in genome size was also reported among landraces from both the northwestern Argentina highlands (NWA; 2C = 4.2–6.3 pg) (Fourastié *et al*. 2018) and the northeastern Argentina lowlands (NEA; 2C = 4.62–6.29 pg) (Realini *et al*. 2016b).

Genome size, when combined with cytogenetic data, can be used to assess differences among landraces, validate groups identified through phenotypic and genetic characterization, and infer the intraspecific dynamics of the maize genome. However, despite the considerable amount of genome size and cytogenetic information available for maize landraces from northeastern Argentina, such information remains limited for maize landraces from the lowland regions of Uruguay and Brazil.

Therefore, this study aimed to evaluate intragenomic variability among maize landraces from Uruguay and Brazil by addressing three fundamental questions: (1) Is genome size variation predominantly structured at the race, landrace, or regional level? (2) Does the number of chromosomal knobs exhibit a similar pattern of structuring and/or reflect regional differentiation? (3) To what extent is genome size variation associated with differences in knob number or other genomic components? Addressing these questions may provide new insights into the history and evolutionary dynamics of the maize genome in the South American lowlands.

## Material and Methods

### Plant Material

Plant material was obtained from the Germplasm Bank of the Universidad de la República (Uruguay). Sampling was selected based on the updated published study reclassifying lowland South American races of Brazil and Uruguay (Silva *et al*. 2020). In total, we evaluated 15 maize landraces representing 13 races, including eight from Uruguay and seven from Brazil (Table S1). Landraces were selected based on information regarding their sequencing and geographic representativeness.

## Methods

### Chromosome preparations and fluorescent hybridization in situ (FISH)

For chromosome preparations, root tips were obtained from seeds germinated in Petri dishes and pretreated with 2 mM 8-hydroxyquinoline for 2 h and 45 min in a dry bath at 25°C, fixed in Carnoy’s solution (3:1 ethanol acid) for at least 24 h at room temperature, and then stored at −20°C. Slides were prepared using the cell dissociation and air-drying technique following the protocol described by Carvalho et al. (2017).

The 180-bp repetitive DNA sequence associated with maize knobs (Peacock *et al*. 1981) was used as a target for probe design to detect knob regions on the chromosomes of maize landraces. A 60-nt DNA oligonucleotide derived from the 180-bp knob repeat was synthesized by Macrogen Inc. (Seoul, South Korea) and labeled with Cy3 at the 5′ end. The pretreatment and in situ hybridization procedures were based on Carvalho et al. (2022) with some modifications. For each chromosome preparation, 20 μL of a hybridization mixture containing the probe was used. The hybridization mixture was denatured at 80°C for 10 min, cooled on ice, applied to the chromosome preparations, and covered with a coverslip. The preparations were then denatured on a heating plate at 80°C for 10 min. Hybridization was performed at 37°C for 16 h. Following hybridization, slides were washed once in 2× saline-sodium citrate (SSC) buffer at room temperature for 5 min, twice in 2× SSC at 42°C for 5 min, twice in 0.1× SSC at 42°C for 5 min, once in 2× SSC at 42°C for 5 min, and finally once in 2× SSC at room temperature for 10 min. The slides were counterstained with 0.2 μg/mL 4′,6-diamidino-2- phenylindole (DAPI) and mounted in 5 μL of Vectashield. Images were captured using a Zeiss AxioImager M2 microscope equipped with a Cohu CCD camera. Brightness and contrast were adjusted using Photoshop CS3. Ten individuals per accession were analyzed, and at least three metaphases per individual were examined to determine the number of knobs and the presence of B chromosomes.

### Genome size measurement

Nuclear DNA content (2C values) was estimated following the procedure of Galbraith et al. (1983) with modifications. Nuclei were isolated using Woody Plant Buffer (Loureiro *et al*. 2006) and stained with propidium iodide (final concentration: 50 µg/mL). Samples were analyzed with a Partec CyFlow® Space flow cytometer (Sysmex-Partec GmbH, Münster, Germany), and data were processed using the FloMax® software (Version 2.4d; Sysmex-Partec GmbH, Münster, Germany). *Solanum lycopersicum* Lehm. ‘Stupické Rané’ (2C = 1.96 pg) was used as the internal standard, and only measurements with a coefficient of variation (CV) < 5% were accepted. Nuclear DNA content (2C) was calculated using the equation: sample peak mean/standard peak mean × 2C DNA content of standard (pg).

### Analysis of repetitive DNA families of maize landraces

To identify DNA sequences associated with differences in genome size, repetitive DNA sequences were analyzed. Total genomic DNA was extracted from fresh leaves of five individuals representing the range of genome sizes, following the CTAB protocol described by Lassner et al. (1989). Low-coverage genome sequencing was performed on a DNBseq platform at the Beijing Genomics Institute (BGI; Shenzhen, Guangdong, China), generating approximately 1× genome coverage per sample with 150-bp paired-end reads. All raw sequencing data have been deposited in the National Center for Biotechnology Information (NCBI) BioProject database under accession number PRJNAXXXXX.

For the identification and quantification of repetitive sequences, similarity-based graph clustering was performed using the RepeatExplorer2 (RE2) pipeline. The pipeline is provided by the ELIXIR-CZ project, part of the international ELIXIR infrastructure (https://www.repeatexplorer-elixir.cerit-sc.cz) (Novák *et al*. 2013, 2017, 2020). Raw DNA-seq reads were uploaded to the RE2 Galaxy server, where their quality was assessed using FastQC v0.11.5 (Andrews 2010). Read preprocessing was performed using the FASTQ paired-end reads tool available in RepeatExplorer Utilities. This tool performed quality filtering using a Phred score cutoff of 20, retaining only reads in which at least 90% of bases exceeded this threshold. The preprocessing tool also converted sequences to FASTA format and interlaced forward and reverse reads into a single file suitable for repetitive sequence analysis (Novák *et al*. 2020). Repetitive sequence analysis was performed using the comparative analysis option implemented in RepeatExplorer2. For this purpose, each file was assigned a prefix, and the files were concatenated into a single file before clustering. Clustering was performed using the default settings of 90% similarity over a minimum sequence overlap of 55%, with sampled paired-end reads equivalent to 0.1× genome coverage. Automated RE2 annotations of clusters representing at least 0.01% of the genome were manually checked and corrected when necessary. The relative abundance of each repeat was calculated based on the number of reads in the corresponding cluster for each accession, excluding organellar reads. The consensus monomer sequences of individual satellite DNA families were subjected to BLASTn searches against their respective genomic assemblies or raw sequencing reads.

### Statistical Analysis

In this study, the 13 maize races, originally classified based on morphological descriptors and named according to country-specific nomenclature, were grouped into four categories (Blanco Dentado, Complejo Cateto, Complejo Moroti-Caingang, and Pisingallo) based on endosperm type to facilitate statistical comparisons between Brazil and Uruguay. The data were divided into two subsets: one comprising genome size measurements from all landraces from Uruguay and Brazil (n = 269 individuals) and the other comprising genome size measurements and knob counts (n = 114 individuals). These variables were compared across countries, races, landraces, grain types (floury, flint, dent, and popcorn), biomes (Caatinga, Atlantic Forest, Pampa–Atlantic Forest ecotone, and Pampa), and latitude. The presence of B chromosomes was also assessed in the sampled individuals. The data were analyzed using RStudio software (version 2026.01.1+403), and figures were generated using *ggplot2*.

Genome size (pg) was analyzed using linear mixed-effects models (LMMs; *lmer* function, *lme4* package), with **Satterthwaite’s approximation used to estimate degrees of freedom** (*lmerTest* package). To test for differences among races and grain types, separate linear mixed-effects models were fitted, with race or grain type as a fixed effect and accession nested within race as a random effect. The general model structure was **Genome size ∼ factor + (1 | Race)**. The significance of fixed effects was assessed using ANOVA, and estimated marginal means and pairwise comparisons were obtained using the *emmeans* package. To assess potential geographic effects, mixed-effects models were fitted with country and its interaction with race or grain type. These models allowed us to test whether differences in genome size between countries depended on race or grain type. Comparisons between countries within each race or grain type were performed using estimated marginal means. To investigate a possible ecological gradient between Brazil and Uruguay, accessions were classified into three regions: the Uruguayan Pampa, the Pampa–Atlantic Forest ecotone in Brazil, and other Brazilian biomes. A mixed-effects model was fitted with region as a fixed effect and accession nested within race as a random effect. Pairwise comparisons between regions were performed using Tukey-adjusted tests. Furthermore, the relationship between latitude and genome size was evaluated using a linear mixed-effects model, with absolute latitude as a continuous predictor and accession nested within race as a random effect. Variation among accessions within each race was evaluated using separate linear models for each race, with accession as an explanatory variable. When appropriate, pairwise comparisons between accessions were performed using Tukey’s adjustment. Finally, the relative contributions of different hierarchical levels to variation in genome size were estimated using a mixed-effects model with random intercepts for country, race nested within country, and accession nested within race. Variance components were expressed as percentages of the total variance.

The number of heterochromatic knobs was analyzed as a count variable and therefore modeled primarily using generalized linear mixed-effects models (GLMMs) with a Poisson distribution and log link function. Initially, the distribution of the response variable, the relationship between the mean and variance, the frequency of zeros, and the presence of overdispersion were evaluated. No evidence of overdispersion was detected. Due to perfect collinearity between country and biome, as well as between race and grain type, only country and race were retained as explanatory variables in the main models. Differences in knob numbers between countries and races were evaluated using generalized linear mixed-effects models, with accession as a random effect. Hierarchical variation among races and among accessions within races was assessed using models with nested random intercepts. Additionally, separate models were fitted for each race to test for differences among accessions, using Poisson models and type II ANOVA. When appropriate, estimated marginal means and Tukey-adjusted pairwise comparisons were obtained using the *emmeans* package. To assess whether patterns of variation in knob number differed between countries, separate models were fitted for Brazil and Uruguay, in addition to a global model including the interaction between country and race. **The significance of this interaction** was tested by comparing nested models with and without the interaction term. Adjusted means for each country–race combination were estimated on the response scale and used for interpretation and graphical representation. The relationship between knob counts and genome size was investigated using simple and multiple linear regression models. Models with and without **knob number as a predictor** were compared, separately controlling for the effects of race, country, and the country × race interaction. A linear mixed-effects model including a random intercept for accessions nested within races was also fitted to account for the hierarchical structure of the data. Associations between latitude, genome size, and knob counts were also evaluated. Latitude was standardized before analysis. The relationship between genome size and latitude was tested using a linear mixed-effects model, whereas the relationship between knob counts and latitude was evaluated using a Poisson generalized linear mixed-effects model, both including accession as a random effect.

## Results

### Intraspecific variation in maize genome size in the lowlands of South America

For maize genome size analysis, a total of 269 individuals were analyzed, 134 from Brazil and 135 from Uruguay, distributed across 15 landraces and classified into four races (Table S1). The mean genome size was 2C = 5.31 pg (5203.8 Mb), ranging from 2C = 4.85 pg (4753 Mb) to 2C = 5.78 pg (5664.4 Mb), representing a range equivalent to 17.5% of the mean. In general, Brazilian races had larger genome sizes than Uruguayan races, although the difference was not statistically significant (Figure 1, Table S2). The adjusted mean genome sizes were 2C = 5.37 pg (5262.6 Mb) for Brazilian races and 2C = 5.27 pg (5164.6 Mb) for Uruguayan races. The same pattern was observed when grain type was analyzed across countries (Figure S1).

**Figure 1.**
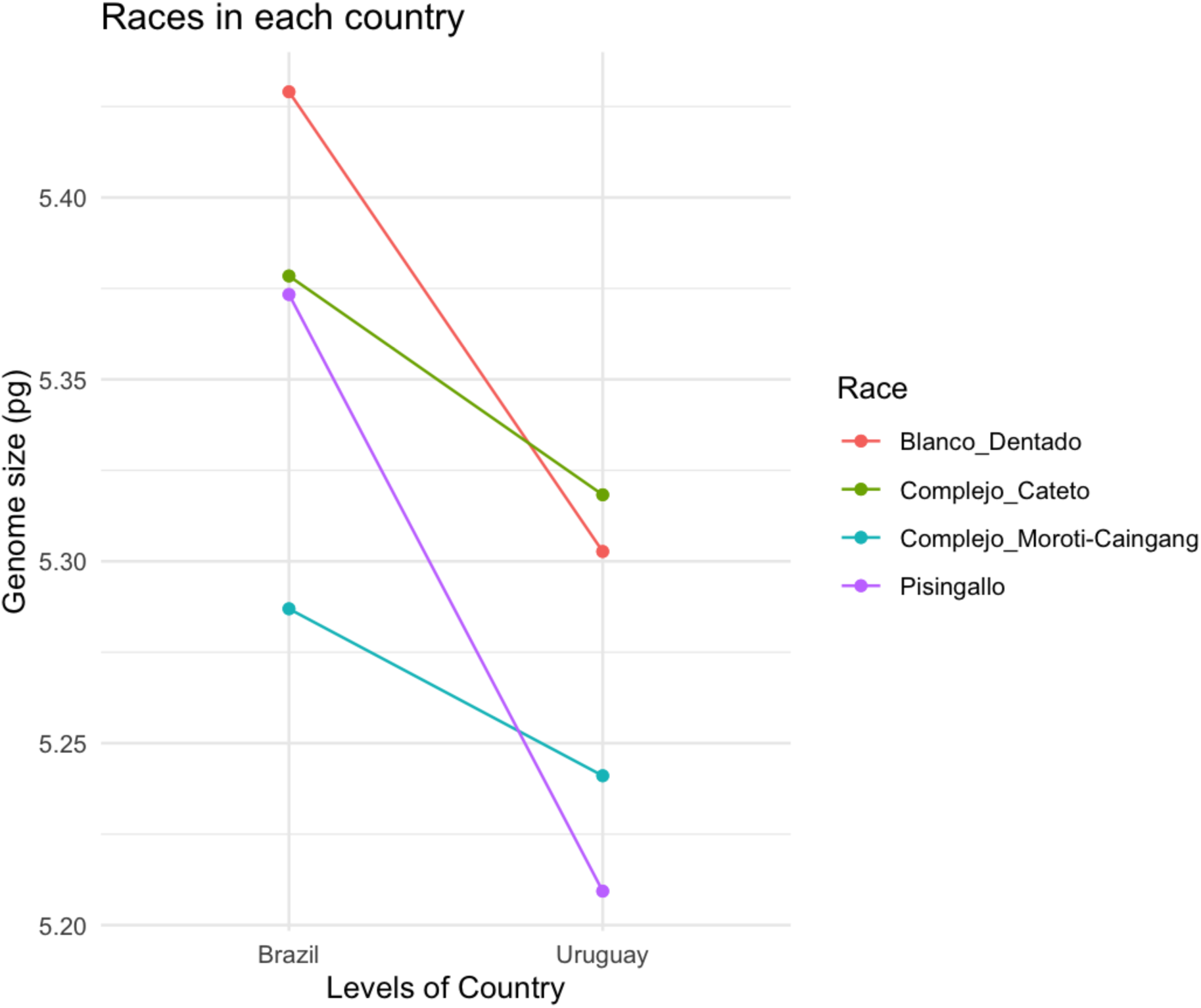
Genome size (2C values) comparisons between Uruguay and Brazil across maize races. Each colored line represents a different maize race.

Significant differences in 2C DNA content (p < 0.05) were detected among landraces (Table 1). Hierarchical analysis of genome size variability and intraracial analysis showed that landraces accounted for most of the intraspecific variation in maize genome size (Figure 2 and Figure S2).

**Figure 2.**
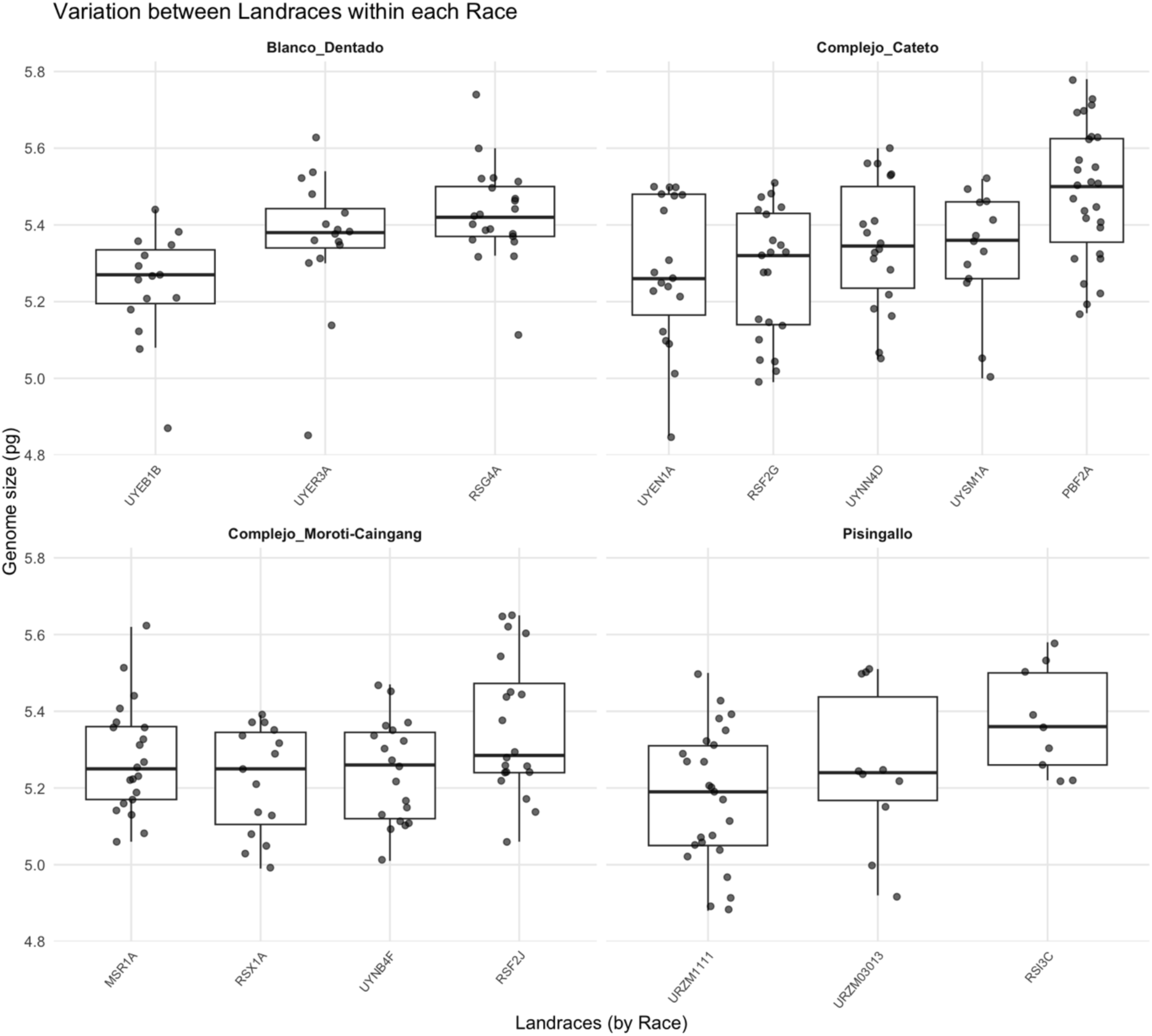
Genome size variation among maize landraces within each race. Boxplots show the distribution of genome size across the accessions analyzed for each race. Genome size is expressed in picograms (pg) on the y-axis, and the x-axis shows the accessions within each race. UR, UY - Uruguay; PB, MS, RS - Brazil.

**Table 1.**
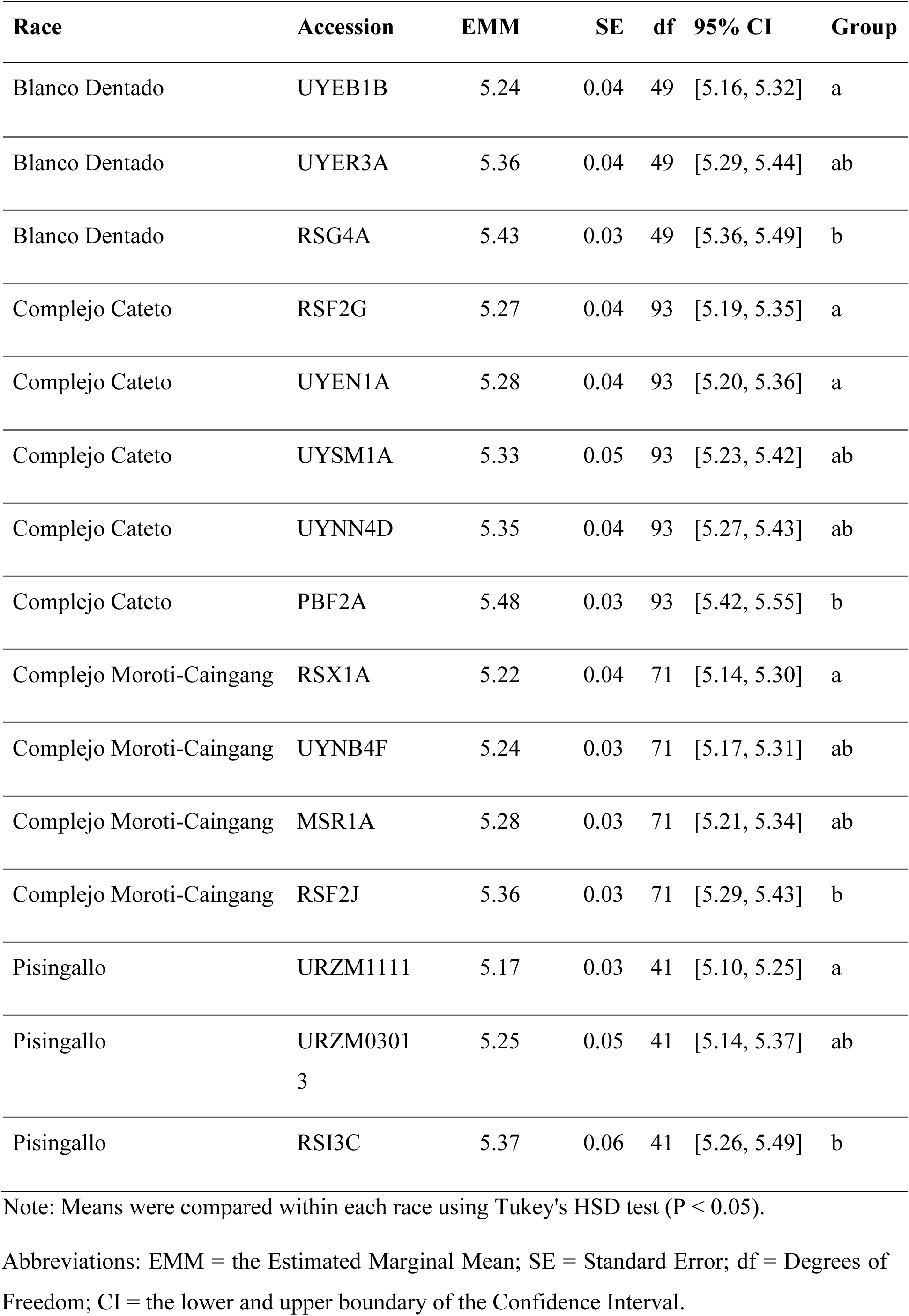
Variation in genome size among accessions within each race.

For analyses involving biomes, the landraces were reorganized into three groups to test whether landraces from the Atlantic Forest–Pampa Ecotone exhibited intermediate genome sizes between those from Brazilian biomes and the Uruguayan Pampa. Landraces cultivated in the Caatinga and Atlantic Forest were grouped as Other Brazilian biomes, landraces cultivated in the state of Rio Grande do Sul were grouped as Ecotone-RS, and landraces from Uruguay were grouped as Pampa-Uruguay. No significant differences in genome size were found among groups, although a decreasing trend in genome size from north to south was observed (Figure S3, F_2, 12.04_= 1.55, p = 0.250). This trend was also evaluated using latitude. In this analysis, genome size was significantly associated with latitude, consistent with a decreasing gradient in landrace DNA content from northeastern Brazil to Uruguay (north to south (Figure 3, F_1, 11.41_ = 7.11, p = 0.020).

**Figure 3.**
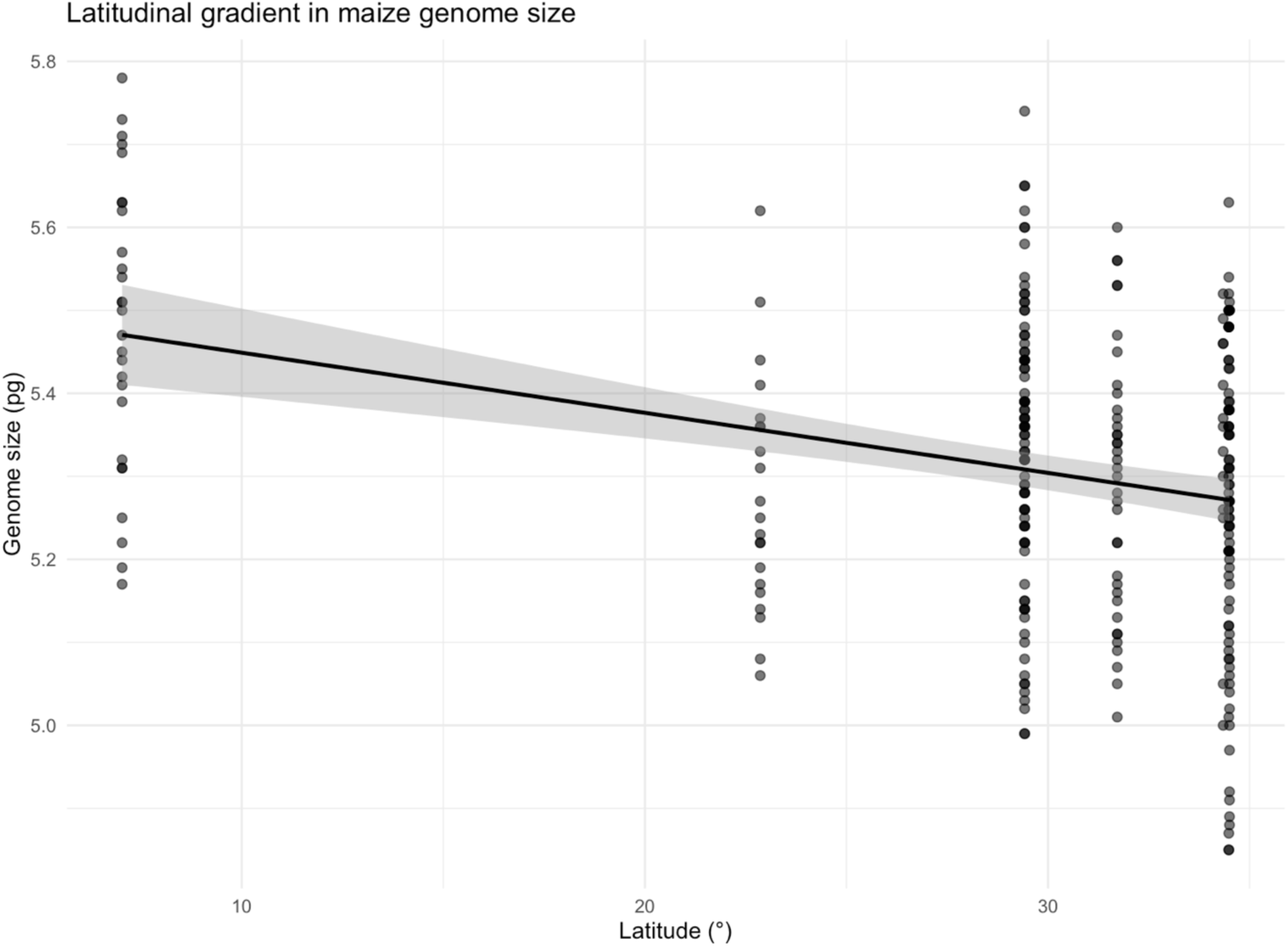
Latitudinal gradient in maize genome size (F_1, 11.41_ = 7.11, p = 0.020). Latitude on the x-axis is in absolute value and genome size in picograms on the y-axis.

### Cytogenetic and statistical characterization of knob variation in maize landraces from the lowlands of Uruguay and Brazil

Analysis of 114 individuals using fluorescent in situ hybridization (FISH) with the 180- bp knob repeat probe revealed variation in the number of heterochromatic knobs among maize landraces, with counts ranging from 13 to 39 knobs per individual. Variation was also observed in knob size and chromosomal position (Figure 4).

**Figure 4.**
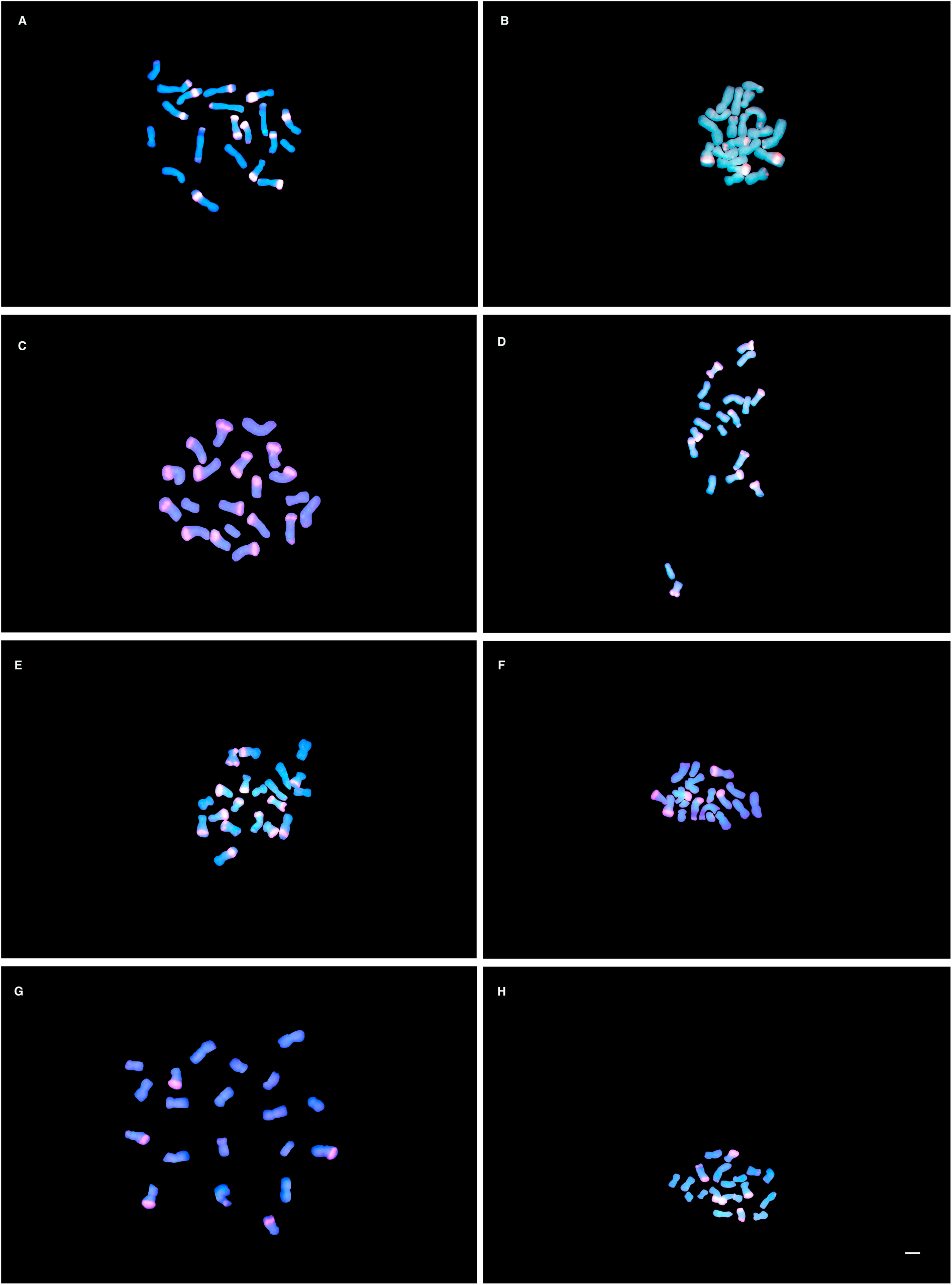
FISH mapping of intragenomic variation of knob number in maize races from Brazil and Uruguay, using a probe for the 180-bp knob repeat (red). A. Complejo Cateto race from Brazil - 26 knobs, B. Complejo Cateto race from Uruguay - 17 knobs, C. Dentado race from Brazil - 31 knobs, Dentado race from Uruguay - 16 knobs, E. Pisingallo race from Brazil - 23 knobs, F. Pisingallo race from Uruguay - 28 knobs, G. Complejo Moroti-Caingang race from Brazil - 19 knobs, H. Complejo Moroti-Caingang race from Uruguay - 27 knobs. Scale bar = 10 um

In the overall analyses of knob number, no significant differences were detected between countries or among races. However, when separate models were fitted for each race to test for differences among accessions within races, significant differences in knob number were detected only in Blanco Dentado (χ² = 16.88, df = 2, p < 0.001; Table S3). Tukey’s post hoc comparisons revealed significant differences among accessions from Brazil and Uruguay. Furthermore, a model testing whether the pattern of variation in knob number among races differed between countries revealed a significant country × race interaction (p < 0.001), indicating that the effect of race on knob number depended on the country (Table 2).

**Table 2.** Fixed effects of the generalized linear mixed model predicting the pattern of variation in the number of knobs according to maize race and country of origin.

| Term/Predictor | Estimate ($\beta$) | SE | Statistic | p-value |
| --- | --- | --- | --- | --- |
| (Intercept) | 3.34 | 0.08 | 43.60 | < 0.001 |
| Country |  |  |  |  |
| Uruguay | -0.38 | 0.10 | -4.01 | < 0.001 |
| Race |  |  |  |  |
| Complejo Cateto | -0.11 | 0.09 | -1.17 | 0.241 |
| Complejo Moroti-Caingang | -0.26 | 0.09 | -2.99 | 0.003 |
| Pisingallo | -0.21 | 0.10 | -2.06 | 0.039 |
| Interactions (Country × Race) | | | | |
| Uruguay x Complejo Cateto | 0.26 | 0.12 | 2.24 | 0.025 |
| Uruguay x Complejo Moroti-Caingang | 0.45 | 0.13 | 3.36 | 0.001 |
| Uruguay x Pisingallo | 0.43 | 0.13 | 3.37 | 0.001 |

Note: Reference level for Country: Brasil; reference level for Race: Blanco Dentado. Values correspond to the model's untransformed estimates. Abbreviations: SE = Standard Error

First, Poisson generalized linear mixed-effects models fitted separately for each country revealed distinct patterns of variation in knob number. In both countries, the estimated variance among accessions within races was zero. However, races differed significantly in the mean number of heterochromatic knobs in both countries (p < 0.05; Tables S4 and S5). In Brazil, the Pisingallo and Complejo Moroti-Caingang races had fewer knobs than the Blanco Dentado race, whereas in Uruguay, these same races had more knobs than Blanco Dentado.

Second, a model including the country × race interaction revealed a significant interaction, indicating a reversal in the pattern of knob number among races between countries (Table 2). Blanco Dentado and Complejo Cateto exhibited higher knob numbers in Brazil than in Uruguay, whereas Pisingallo and Complejo Moroti-Caingang exhibited higher knob numbers in Uruguay than in Brazil (Figure 5).

**Figure 5.**
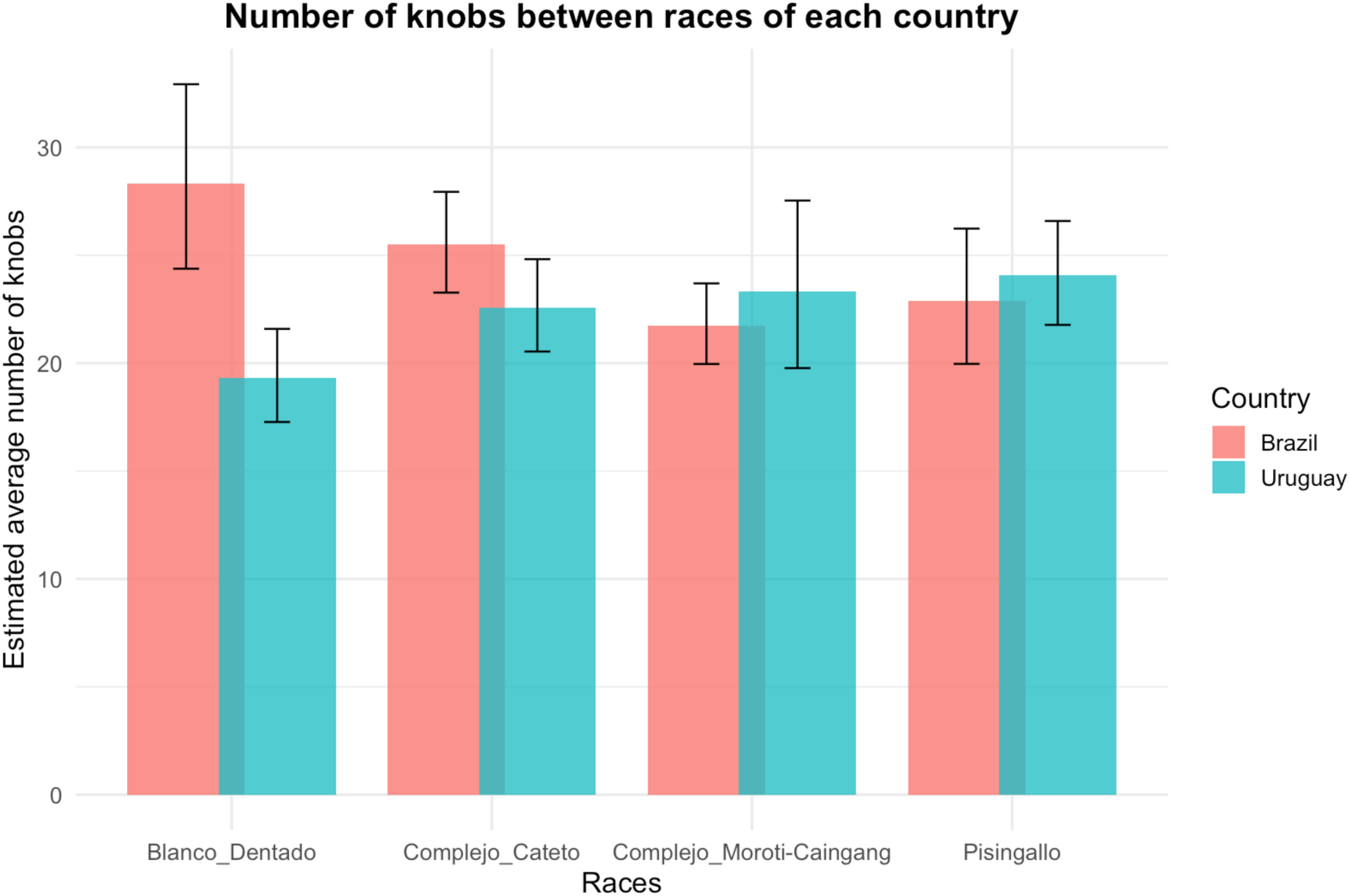
Knob numbers among races in each country. Adjusted means were estimated on the knob number scale (y-axis), and the vertical error bars indicate the 95% asymptotic confidence intervals (x-axis).

### Relationship between knobs number and genome size

The data from this subset (comprising genome size and knob number measurements for 114 individuals) were analyzed using models ranging from simple linear regression to linear mixed-effects models accounting for the hierarchical structure of the data. All models consistently showed no significant association between knob number and genome size (Figure S4 and Table S6).

The latitudinal gradient in genome size remained significant in the subset of individuals with knob data (F_1, 9.77_ = 5.63, p = 0.040), consistent with the decrease in genome size from northeastern Brazil to Uruguay observed in the full dataset. However, knob number was not significantly associated with latitude and did not significantly explain variation in genome size, suggesting that the observed cline in DNA content may not be directly related to variation in the number of these heterochromatic regions (Tables S7 and S8).

### Analysis of repetitive content in maize landraces

The landraces selected for these analyses were representative of the north–south gradient of decreasing genome size. The PBF2A landrace (Caatinga biome) had the largest genome size, followed by RSG4A (Atlantic Forest–Pampa ecotone), both from Brazil, whereas the UYNN4D landrace (Pampa biome) had the smallest genome size. The repetitive DNA fraction of the five genomes analyzed ranged from 1,856.92 Mbp (70.83%) to 1,922.63 Mbp (72.31%). The proportions of annotated repetitive DNA are presented in Table 3 and Figure 6.

**Figure 6.**
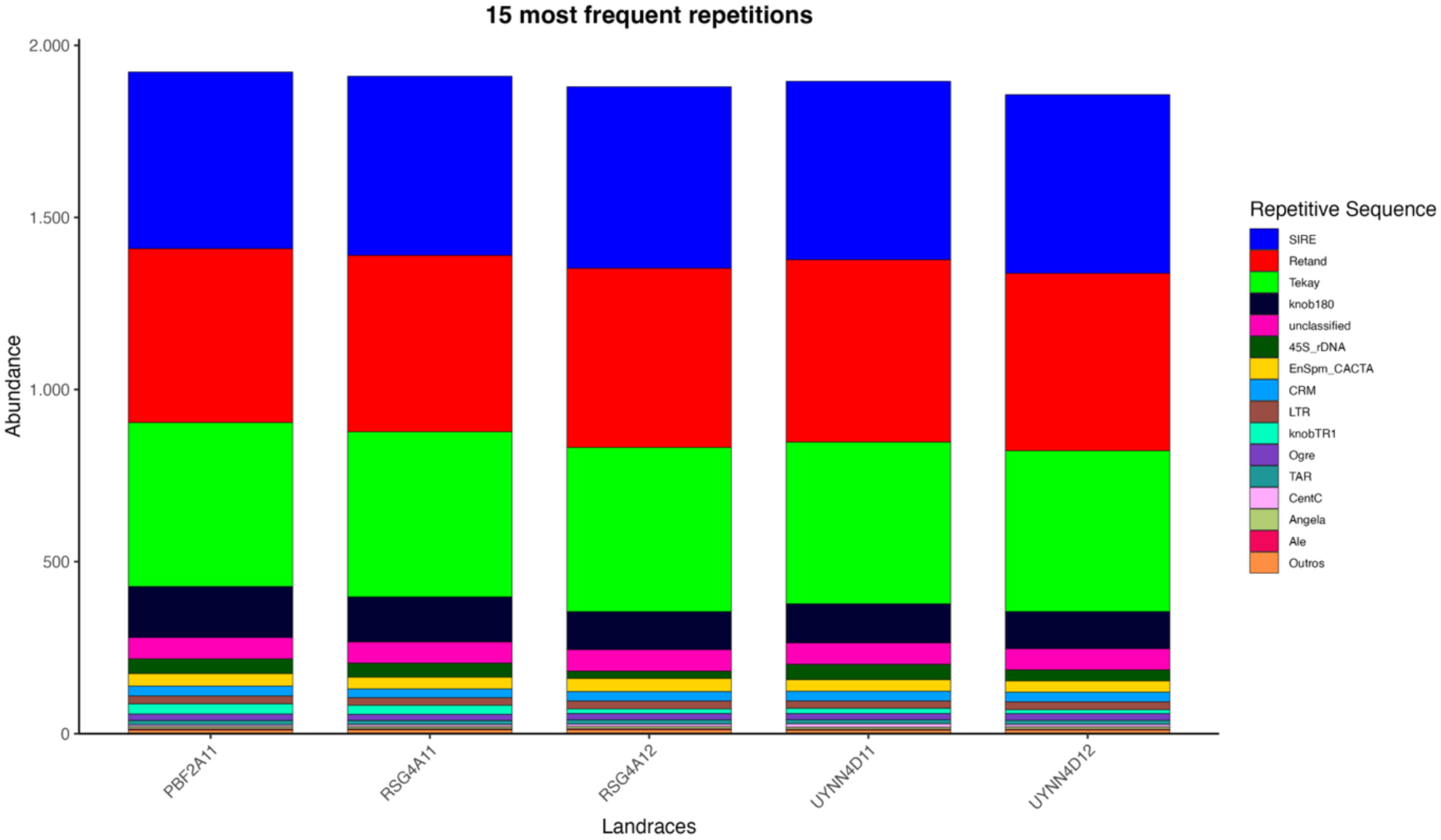
Abundance (in Mb) of the most common repetitive DNA elements in the genomes of maize landraces.

**Table 3.** Genomic proportions (in Mb) of repeat classes and families of maize landraces from Brazil and Uruguay.

| **Repeat Classification** | **PBF2A11**<br>**(Mpb)** | **RSG4A11**<br>**(Mpb)** | **RSG4A12**<br>**(Mpb)** | **UYNN4D12**<br>**(Mpb)** | **UYNN4D11**<br>**(Mpb)** |
| --- | --- | --- | --- | --- | --- |
| All/repeat/mobile_element/Class_I/LINE | 1.44 | 1.08 | 1.32 | 1.17 | 1.10 |
| All/repeat/mobile_element/Class_I/LTR/Ty1_copia | 538.86 | 544.51 | 553.44 | 542.37 | 543.63 |
| All/repeat/mobile_element/Class_I/LTR/Ty3_gypsy | 1029.85 | 1036.59 | 1043.81 | 1046.68 | 1030.76 |
| All/repeat/mobile_element/Class_II/Subclass_1/TIR | 37.03 | 35.25 | 39.63 | 35.25 | 33.75 |
| All/repeat/mobile_element/Class_II/Subclass_2/Helitron | 1.05 | 1.43 | 1.34 | 1.21 | 1.35 |
| All/repeat/rDNA/45S_rDNA | 43.42 | 41.29 | 21.40 | 45.00 | 32.63 |
| All/repeat/rDNA/5S_rDNA | 1.04 | 0.96 | 0.59 | 0.67 | 1.14 |
| All/repeat/satellite | 185.36 | 166.13 | 132.85 | 140.40 | 128.93 |
| unclassified | 61.62 | 61.11 | 62.26 | 61.74 | 60.81 |

Among transposable elements, long terminal repeat (LTR) retrotransposons are the most abundant repetitive elements in the maize landrace genomes (58.42%–60.61%), with the Ty3-gypsy and Ty1-copia superfamilies being the most frequent, followed by satellite DNA sequences (Figure 7). The Retand lineage was the most abundant among Ty3-Gypsy elements, followed by Tekay (Table S9), whereas the SIRE lineage was the most abundant among Ty1-Copia elements. DNA transposons accounted for less than 2% of the genomes analyzed, with the CACTA lineage being the most abundant in all landraces.

**Figure 7.**
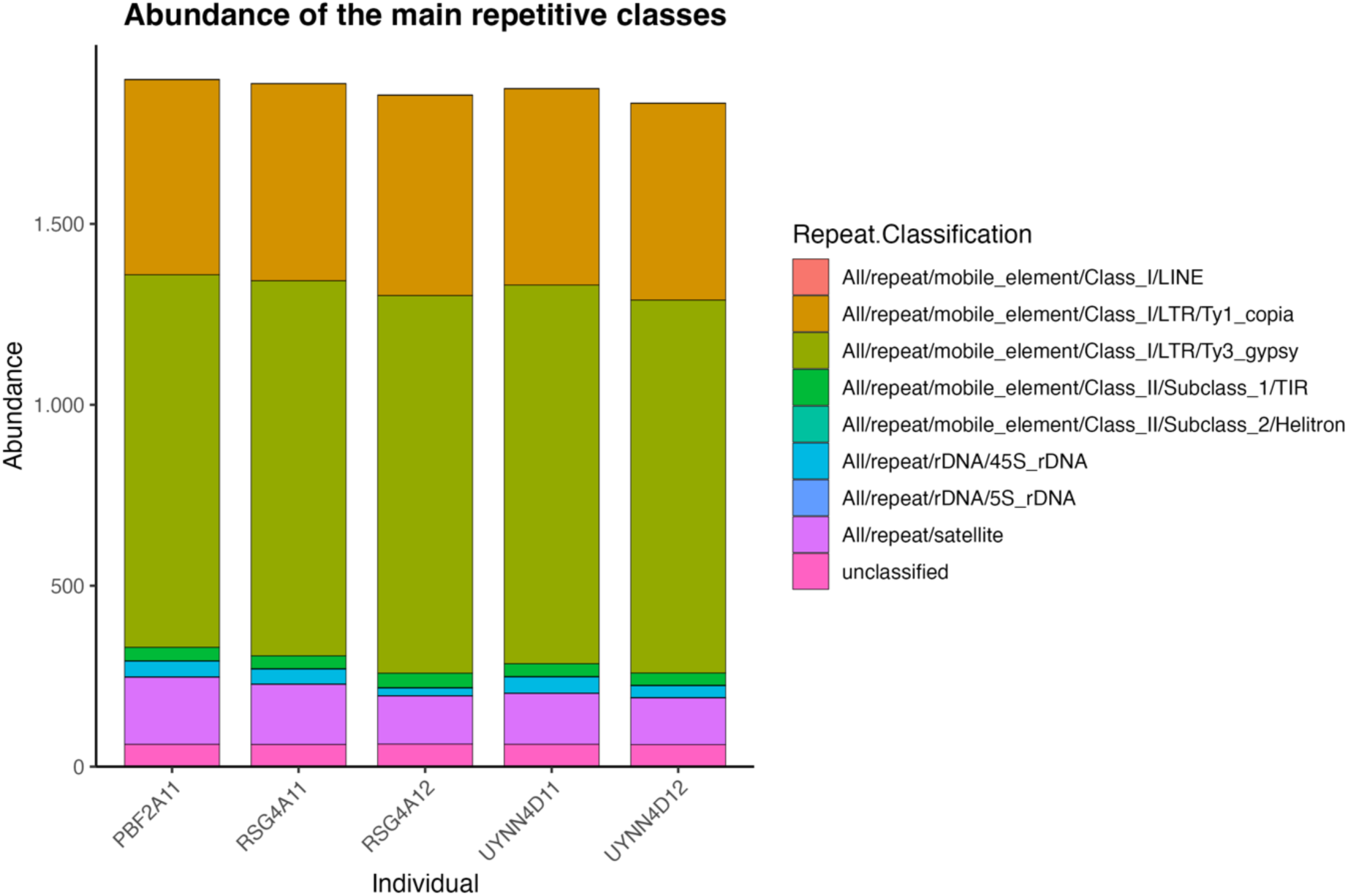
Repetitive DNA genome sequences in maize landraces, showing the abundance (in Mb) of LINEs, Ty1-Copia and Ty3-Gypsy retrotransposons, TIR and Helitron transposons, rDNA (45S and 5S), satellite DNA, and unclassified repetitive sequences.

Satellite DNA sequences accounted for 4.91%–6.90% of the genomes. Among tandem repeat sequences, the 180-bp knob repeat was the most abundant satellite DNA family. In addition, CentC and Cent4 satellite sequences, the 531-bp telomeric sequence, and the 350-bp TR-1 knob repeat, which was much less abundant than the 180-bp knob repeat, were identified (Figure 8). Ribosomal DNA sequences were also detected.

**Figure 8.**
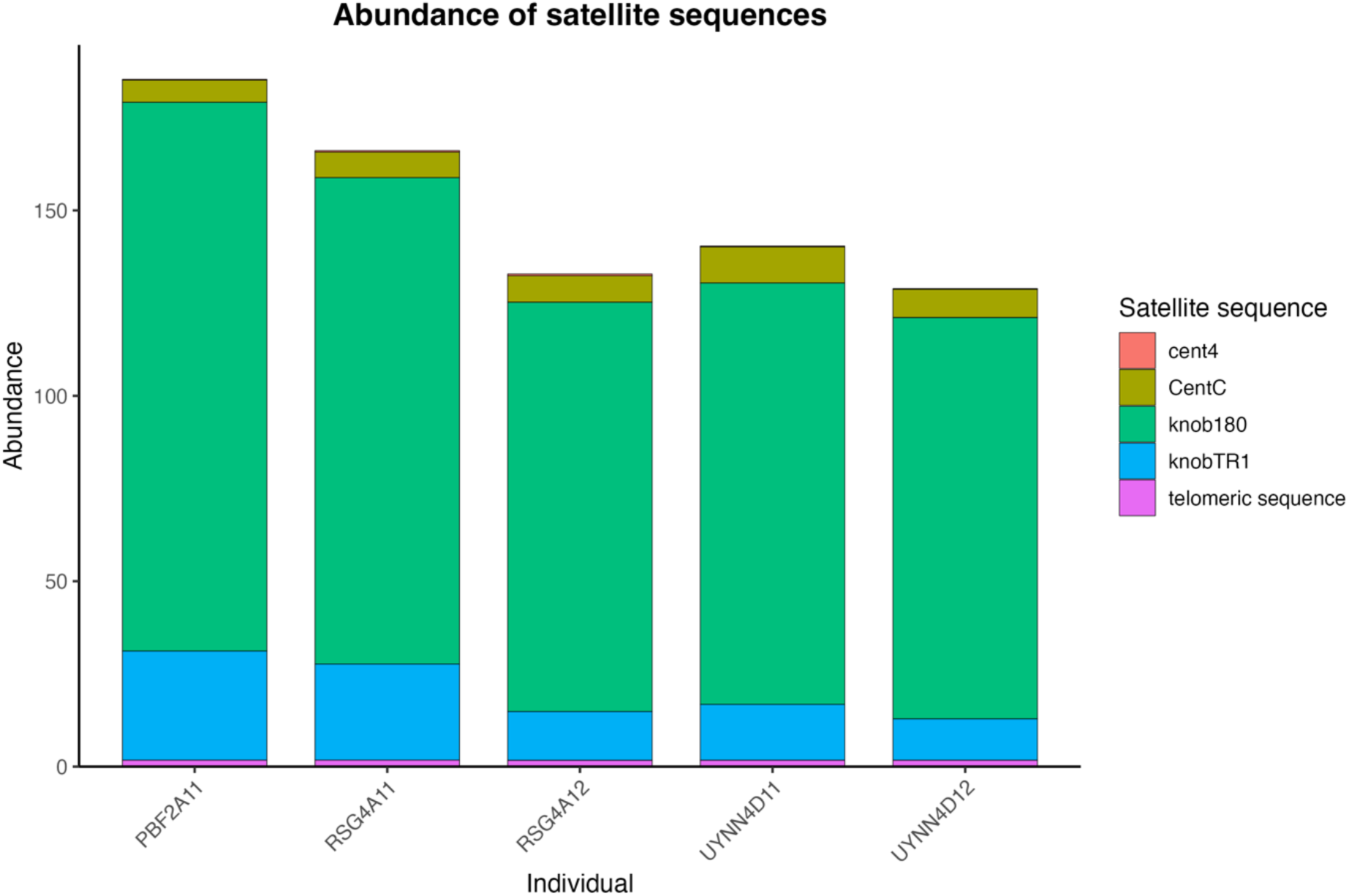
Genomic abundance (in Mb) of satellite DNA families in maize landraces.

A trend toward increasing retrotransposon abundance was observed with decreasing genome size among landraces, whereas satellite DNA abundance tended to decrease. When repetitive element families were analyzed in greater detail, the 180-bp and TR-1 satellite sequences showed a decreasing trend, whereas the SIRE and Retand lineages showed an increasing trend and were the most abundant retrotransposon lineages.

### B chromosomes analysis

Finally, B chromosomes were detected in Uruguayan landraces but not in any of the Brazilian landraces analyzed. In total, B chromosomes were detected in three of the eight Uruguayan landraces analyzed, representing two maize races: Pisingallo and Complejo Cateto. The frequency of individuals carrying B chromosomes was approximately 0.44. Polymorphism in B chromosome number, ranging from one to two chromosomes per individual, was also observed within landraces. For example, URZM03013 carried two B chromosomes, whereas UYEN1A carried one (Figure 9; Table S10).

**Figure 9.**
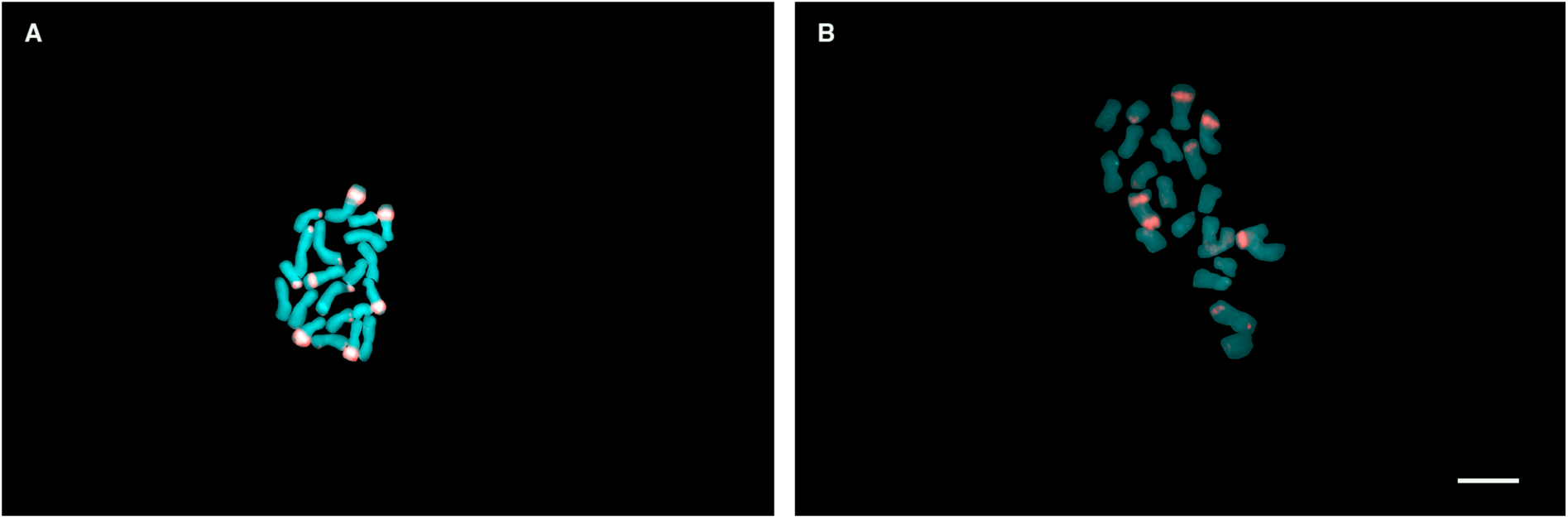
B chromosomes in Uruguayan landraces. A) UYEN1A landrace carrying one B chromosome (2n = 21) and B) URZM03013 landrace carrying two B chromosomes (2n = 22). Scale bar = 10 um

## Discussion

### Latitudinal cline of maize genome size

This study provides the first characterization of DNA content in maize races and landraces from the South American lowlands of Brazil and Uruguay. Variation in maize genome size is recognized as a key component of its intraspecific diversity and has been extensively documented across races and landraces (Laurie and Bennett 1985; Rayburn *et al*. 1985; Poggio *et al*. 1998; Díez *et al*. 2013). In the present study, genome size among landraces ranged from 4.85 pg to 5.78 pg, representing a 17.5% variation. This variation was lower than that reported by Poggio et al. (1998), Realini et al. (2015) and Fourastié et al. (2018) for Argentinian landraces, in which genome size variation ranged from 36% to 42%, corresponding to 2C values between 4.4 and 6.8 pg.

In general, maize races from Brazil had larger genomes than those from Uruguay, although this difference was not statistically significant. This result suggests that variation in genome size may not be abrupt between these countries but rather continuous across geographic space. Differences in DNA content are commonly observed in plants, and two main evolutionary processes contribute to this variation: polyploidy and changes in repetitive DNA sequences (Dodsworth *et al*. 2015). Throughout evolution, these processes can lead to increases or decreases in genome size, resulting in genetic changes, including the gain or loss of genes, that can have substantial effects on the phenotype (Soltis and Soltis 2003; Soltis *et al*. 2015). In maize, this variability has been associated with the presence of heterochromatic knobs, transposable elements, and variation in the number and size of B chromosomes (Laurie and Bennett 1985; Rayburn *et al*. 1985; Rayburn and Auger 1990; Poggio *et al*. 1998; Rosado *et al*. 2009; Díez *et al*. 2013; Tenaillon *et al*. 2016; Fourastié *et al*. 2018; Bilinski *et al*. 2018).

Analyses by biome revealed a consistent pattern of decreasing DNA content from north to south, although the differences were not statistically significant. In contrast, when latitude was incorporated into the analyses, a significant association with genome size was detected, supporting the existence of a north–south latitudinal gradient. Bureš et al. (2024) analyzed the global distribution of genome sizes across more than 16,000 angiosperm species and identified a latitudinal trend in genome size (an S-shaped curve), in which species with smaller genomes occurred in tropical and Arctic regions, whereas species with larger genomes occurred in temperate regions, supporting the large genome constraint hypothesis. In lowland maize, however, we observed an opposite latitudinal trend: landraces closer to the equator had larger genomes, whereas those in temperate regions had smaller genomes.

Laurie and Bennett (1985) and Rayburn et al. (1985) reported that the DNA content of maize from the Northern Hemisphere decreased significantly with increasing latitude. In the present study, a similar pattern was observed in the Southern Hemisphere, with genome size decreasing from north to south. This pattern is also consistent with findings in other species, such as *Rorippa indica* (Han *et al*. 2024), *Crithmum maritimum* (Roxo *et al*. 2023) and *Cynodon dactylon* (Zhang *et al*. 2019). Interestingly, a recent study on the evolutionary history of maize suggested that genome size decreased as populations moved away from the center of domestication and that this process occurred along dispersal gradients during adaptation to new environments (Kistler *et al*. 2020).

In South America, previous studies have shown associations between genome size variation and altitudinal gradients, suggesting that cytogenetic diversity among landraces in the Southern Cone is not randomly distributed but may reflect structured patterns across geographic space and may be associated with the differential dynamics of repetitive DNA sequences (Poggio *et al*. 1998; Fourastié *et al*. 2018; Realini *et al*. 2021).

The recurrence of altitudinal or latitudinal gradients has been interpreted as a result of indirect selection mediated by developmental traits, such as life cycle duration and growth rate. Smaller genomes are often associated with faster cell cycles, which may favor development in environments at higher latitudes, where the growing season is shorter (Bennett 1987; Poggio *et al*. 1998; Bilinski *et al*. 2018). In this context, the observed association between latitude and genome size is consistent with the hypothesis that variation in nuclear DNA content may be influenced by selective pressures associated with local adaptation, although further investigation is needed to test this hypothesis. In summary, these results suggest that variation in maize genome size may follow a gradual pattern, influenced by geographic gradients and diversity among landraces.

### Intragenomic analysis of knob numbers

Heterochromatic knobs are a prominent genomic feature of maize chromosomes (Albert *et al*. 2010), and since their first description in 1929, numerous studies have sought to elucidate their role in the genome (McClintock 1929; McClintock *et al*. 1981). Our results revealed substantial variation in knob number (13 to 39 knobs per individual) among races and within landraces. The cytogenetic diversity of landraces from northeastern and northwestern Argentina has been extensively characterized. In populations from northwestern Argentina, which are cultivated at high altitudes, negative correlations have been reported between altitude and both the percentage of heterochromatin and knob number, with the latter ranging from 1 to 14 (Poggio *et al*. 1998), 5 to 19 (Fourastié *et al*. 2018), and, more recently, 3 to 15 (Realini *et al*. 2026). In landraces from northeastern Argentina (lowlands), knob number ranged from 10 to 22 (Realini *et al*. 2016) and 8 to 23 (Realini *et al*. 2018). Such variation has also been documented in different types of germplasm, including inbred lines and open-pollinated varieties (Rayburn *et al*. 1985; Mondin *et al*. 2014; Carvalho *et al*. 2022). Inbred lines and open-pollinated varieties characterized by Rayburn et al. (1985) exhibited knob numbers ranging from 2 to 16 and 0 to 18, respectively. Inbred lines and hybrids developed to carry specific knob combinations at defined chromosomal positions also exhibited variation, with knob numbers ranging from 12 to 18 (Mondin *et al*. 2014) and 8 to 14 (Carvalho *et al*. 2022). Overall, higher knob numbers were observed in the present study.

Unlike genome size, which exhibited a continuous clinal pattern, knob number showed a significant race × country interaction, characterized by a reversal in the relative ranking of races between Brazil and Uruguay. This pattern suggests that variation in knob number may reflect differences in genetic history and racial structure. These findings are consistent with those of McClintock et al. (1981), who characterized American maize races based on knob characteristics (number, size, and position), B chromosome frequency, and the presence of abnormal chromosome 10, demonstrating that races possess distinct chromosomal identities and geographic structure. The authors attributed this regional variation to dispersal history, local selection, and introgression among races.

The evolutionary history of maize is complex, and recent evidence indicates that modern maize resulted from multiple episodes of human-mediated selection and dispersal across the Americas (Kistler *et al*. 2020). Previous studies have provided evidence for the arrival of partially domesticated maize in southwestern Amazonia, South America, between 7,500 and 7,000 years before present (BP), where these populations underwent divergent processes of selection and adaptation that also continued in Mesoamerica in sympatry with teosinte populations (Kistler *et al*. 2018; Costa *et al*. 2022). Genomic analyses of archaeological remains have revealed evidence of a second wave of maize dispersal, characterized by increased gene flow from northern *Zea mays* populations. Upon reaching South America, these populations were incorporated into Indigenous agricultural systems and hybridized with previously adapted South American populations (Kistler *et al*. 2020; Yang *et al*. 2023). Genomes recovered from archaeological remains at the El Gigante rock shelter (Honduras), dating to between 2,300 and 1,900 BP, provide evidence of another dispersal event and a clear influence of this second wave, involving lineages that had already hybridized with South American populations and subsequently interbred with Pan-American lineages. According to these studies, hybridization with adapted South American populations enhanced hybrid vigor and contributed to the development of more productive maize populations, coinciding with the transition to a maize-based staple diet in Mesoamerican regions (Kistler *et al*. 2020; Yang *et al*. 2023).

McClintock et al. (1981) reported that South American maize races exhibited distinct knob profiles compared with races from the center of origin, Central America, and the United States. These differences may reflect a legacy of the first wave of maize dispersal into South America, highlighting the importance of this gene pool in the evolutionary history of maize and the value of these genomic elements as cytological markers for understanding genetic diversity within maize. Several studies of South American maize have reported significant correlations between knob characteristics and altitudinal clines (Poggio *et al*. 1998), the presence of B chromosomes (Fourastié *et al*. 2018), genome size (Tito *et al*. 1991) and quantitative traits such as flowering time and seed weight (Realini *et al*. 2021; Carvalho *et al*. 2022).

In the lowlands of South America, which are considered a secondary center of genetic diversity for maize (Brieger 1958; Paterniani and Goodman 1977), Costa et al. (2022) proposed an association between maize dispersal routes and endosperm types, suggesting that this process was mediated by different Indigenous groups. Their study identified at least three dispersal routes—northern, southern, and along the Atlantic coast—associated with the migrations of Indigenous groups belonging to the Arawak, Tupi-Guarani, and Macro-Jê linguistic families.

Furthermore, evidence suggests that popcorn and floury maize were the kernel types introduced into South America during the first wave of dispersal, whereas flint maize was introduced during the second wave (Brieger 1958; Paterniani and Goodman 1977; Kistler *et al*. 2018, 2020; Costa *et al*. 2022). Dent maize was the last kernel type to arrive in Brazil, introduced by American immigrants in the late 19th century (Paterniani *et al*. 2000; Silva *et al*. 2020). In the present study, Blanco Dentado (dent kernel type) was the only race that exhibited pronounced differences in knob number among landraces. Interestingly, the Brazilian landrace that differed significantly from the Uruguayan landraces originated from the state of Rio Grande do Sul, which borders Uruguay. Consistent with this finding, the significant race × country interaction indicates that the relative ranking of races according to knob number differs between Brazil and Uruguay. This raises the hypothesis that Uruguayan dent maize landraces may have a different origin and dispersal history from their Brazilian counterparts.

In the context of maize dispersal history, racial classification is a relatively recent concept developed to organize maize diversity based on phenotypic characteristics, with maize races often associated with specific endosperm types. Here, we found a significant race × country interaction, characterized by a reversal in the relative ranking of races according to knob number between Brazil and Uruguay. This result is consistent with the concepts of race and landrace, as races are defined as groups of landraces that share recognizable phenotypic characteristics and are adapted to particular geographic and sociocultural contexts (Anderson and Cutler 1942; Brieger 1958). Landraces, in turn, are genetically dynamic populations maintained by farmers within traditional agricultural systems. These populations can evolve within their original areas of distribution under farmer-mediated and environmental selection while maintaining their genetic identity through continued cultivation and conservation (Villa *et al*. 2005; Costa *et al*. 2017; Guzzon *et al*. 2021). This dynamic is consistent with the pattern observed in the present study, characterized by variation in knob number within landraces and the grouping of this variation into races. Taken together, these results reveal differences in knob profiles among racial groups and countries, which may reflect distinct demographic and dispersal histories. Further population genetic studies are needed to assess the potential contributions of local selection and introgression to these patterns.

### Knob number does not explain variation in maize genome size

Repetitive sequences shape the genomic landscape through structural rearrangements (e.g., illegitimate recombination, deletions, inversions, translocations, and duplications) and can lead to substantial changes in genome size (Šatović-Vukšić *et al*. 2025). In maize, heterochromatic knobs, in particular, have been considered major contributors to the extensive variation in genome size. However, our results did not reveal a relationship between genome size variation and knob number among maize landraces.

This relationship has been investigated in previous studies, some of which reported positive correlations (Rayburn *et al*. 1985; Tito *et al*. 1991; Jian *et al*. 2017; Fourastié *et al*. 2018), whereas others found no such correlation (Laurie and Bennett 1985; Realini *et al*. 2016a). This study provides the first karyotypic characterization of landraces from Brazil and Uruguay. We hypothesize that the association between heterochromatic knobs and variation in total DNA content may be influenced not only on the number of knobs but also on their size and sequence composition, as suggested by our analysis of repetitive DNA content. Future studies should therefore consider these additional characteristics when investigating this relationship.

### Abundance of repetitive sequences in maize landraces

The maize genome is relatively large, dynamic, and complex, with repetitive sequences accounting for over 80% of its content (Schnable *et al*. 2009; Chia *et al*. 2012; Chen *et al*. 2023). Given the range of genome sizes observed in maize, it is important to investigate the genomic components contributing to this variation. Here, we used low-coverage sequencing to explore the repetitive DNA composition of five individuals representative of the latitudinal gradient in genome size and observed substantial variation in the abundance of individual families of transposable elements and satellite DNA. Overall, transposable elements accounted for approximately 60% of the genomes of the analyzed landraces, whereas satellite DNA represented nearly 7%. Consistent with previous studies, Ty3-gypsy and Ty1-copia were the most abundant transposable element superfamilies identified in our analysis (Hufford *et al*. 2021; Ou *et al*. 2024).

Tenaillon et al. (2011) compared the transposable element content of maize and *Zea luxurians*, showing that these elements account for approximately 70% of the difference in genome size between the two species. At the intraspecific level, Ou et al. (2024) generated a transposable element annotation for a maize pangenome derived from 26 inbred lines that served as founders of the maize nested association mapping population. Their study revealed an excess of 35.1 Mb of transposable element sequences per genome in tropical maize compared with temperate maize, with LTR retrotransposons driving these differences.

Although previous studies have reported that approximately 85% of the maize genome consists of transposable elements (Schnable *et al*. 2009; Chia *et al*. 2012; Chen *et al*. 2023), we found a lower proportion of these elements (60–62%). We hypothesize that this discrepancy may result from methodological differences in the analysis of repetitive sequences or reflect a genuinely lower abundance of transposable elements in maize landraces from the South American lowlands compared with maize from other regions of the continent. However, Tenaillon et al. (2011) showed that even small samples ranging from 1.5 to 5 million bp do not affect estimates of the relative abundance of transposable element families, using an approach similar to that implemented in RepeatExplorer.

Knobs can also vary in their molecular composition among chromosomes and landraces. They are primarily composed of 180-bp repeats and TR-1 elements, and individual knobs may consist exclusively of either repeat type or contain a mixture of both (Ananiev *et al*. 1998; Kato *et al*. 2004; Albert *et al*. 2010; Ghaffari *et al*. 2013; Mondin *et al*. 2014). Here, we physically mapped only the 180-bp knob sequences, but our sequencing data also allowed us to identify TR-1 repeats, which were less abundant than 180-bp repeats. This finding is consistent with previous studies (Realini *et al*. 2016a; Bilinski *et al*. 2018; Chen *et al*. 2023).

A trend toward higher retrotransposon abundance was observed in landraces with smaller genomes, whereas satellite DNA abundance tended to decrease with decreasing genome size. At the family level, the 180-bp and TR-1 satellite repeats followed this decreasing trend, whereas the SIRE and Retand lineages showed the opposite pattern. These trends are consistent with the findings of Chia et al. (2012) who characterized the maize HapMap and investigated the genomic factors influencing genome size variation by comparing maize lines and teosintes. Their study revealed a negative correlation between transposable element abundance and genome size and suggested that this variation was associated with knob content.

Although we found no relationship between knob number and genome size, the patterns observed in the analysis of knob number among races within each country were consistent with those obtained from the analysis of knob repeat abundance among landraces. PBF2A and UYNN4D belong to the Complejo Cateto race, and our analysis showed that this race had a greater number of knobs in Brazil than in Uruguay. This pattern was consistent with the repetitive DNA analysis, in which the Brazilian individual PBF2A exhibited a higher abundance of knob-associated repeats (180-bp and TR-1) than the Uruguayan individual UYNN4D. Therefore, further studies including a larger number of individuals are needed to confirm our observations regarding the dynamics of maize genome size and its relationship with repetitive sequences.

### B chromosomes detected in Uruguayan landraces

B chromosomes are supernumerary and non-essential chromosomes that are present in some individuals or populations within a species and exhibit irregular, non-Mendelian inheritance (Randolph 1941; Johnson Pokorná and Reifová 2021). In maize, these chromosomes are maintained in populations through mitotic nondisjunction during the second pollen division, producing two sperm cells with unequal numbers of B chromosomes, with the sperm cell carrying B chromosomes preferentially fertilizing the egg (Blavet *et al*. 2021).

McClintock et al. (1981) distinguished between Andean and lowland maize races of South America based on the presence of B chromosomes. These chromosomes were observed at high frequencies in Andean regions, whereas in the lowlands, they occurred at very low frequencies and were absent from most of the analyzed materials. Furthermore, B chromosomes have been studied in populations from northwestern Argentina (Poggio *et al*. 1998; Rosato *et al*. 1998; Fourastié *et al*. 2018). In these studies, the authors examined landraces distributed across a wide range of environments and altitudes. A common finding among these studies was an association between the presence of B chromosomes and altitude among Argentina landraces.

In the present study, B chromosomes were detected exclusively in Uruguayan landraces. Although Uruguay is located in the South American lowlands, B chromosomes have been reported in five Uruguayan races and across all four grain types, including popcorn, which is considered ancestral (Garcia da Rosa 2024). The occurrence of these supernumerary chromosomes in Uruguay, together with previous reports from the country, highlights geographic variation in the distribution of B chromosomes among South American lowland maize populations. Whether this pattern reflects differences in the origins or dispersal histories of these landraces remains to be investigated.

### Conclusion

Taken together, our results suggest that although genome size was significantly associated with latitude, variation in knob number alone does not explain the observed variation in DNA content. Thus, the observed gradient in genome size may reflect the combined effects of multiple repetitive sequences or structural variations not captured by knob counting, reinforcing the multifactorial nature of genome size evolution in *Zea mays*. This distinction between a potentially adaptive signal in DNA content and a structural pattern in knob number associated with racial history highlights the complexity of the processes shaping the genomic architecture of maize along geographic gradients.

## Supplementary Information

Supplementary data are available at Annals of Botany online and consist of the following. Table S1: Raw data on maize landraces from the lowlands of South America. Table S2: ANOVA results for the interaction between race and country. Figure S1: Genome size (2C values) comparisons between Uruguay and Brazil across maize grain type. Figure S2: Components of genome size variance. Figure S3: Ecological gradient of genome size between Brazil and Uruguay. Table S3: Pairwise contrasts of the Blanco Dentado race. Table S4: Fixed effects from the generalized linear mixed model that predicts the pattern of variation in the number of knobs, adjusted for Brazil. Table S5: Fixed effects from the generalized linear mixed model that predicts the pattern of variation in the number of knobs, adjusted for Uruguay. Figure S3: Linear relationship between the knob number and genome size. Table S6: Summary of linear and linear mixed-effects models evaluating the relationship between knob numbers and genome size under different covariate controls. Table S7: Fixed effect from the generalized linear mixed model (GLMM) predicting the relationship between knob numbers and latitude. Table S8: Fixed effect from the generalized linear mixed model (GLMM) predicting the relationship between knob numbers, latitude and genome size. Table S9: Genomic proportions (in Mb) of each repeat class and families of maize landraces from Brazil and Uruguay annotated by Repeat Explorer. Table S10: Races and landraces exhibiting variation in the number of B chromosomes.

## Acknowledgements

R. F. C thanks the National Agency for Research and Innovation for a post-doctoral (PD_NAC_2021_1_168393) scholarship.

## Data availability

All data supporting the conclusions of this study are available in the article and its Supplementary Information.

## AI Assistance

Artificial intelligence was used to assist in drafting the statistical analysis methodology section, reviewing the English text, and generating the code used for the statistical analyses.

