## Supplementary material for "Unveiling the intraspecific genomic diversity of South American lowland maize landraces": Suplementary Information

**Table of Contents:**

| Table S2 | Page 2 |
| --- | --- |
| Figure S1 | Page 2 |
| Figure S2 | Page 3 |
| Figure S3 | Page 4 |
| Table S3 | Page 4 |
| Table S4 | Page 5 |
| Table S5 | Page 5 |
| Figure S4 | Page 6 |
| Table S6 | Page 7 |
| Table S7 | Page 8 |
| Table S8 | Page 8 |
| Table S9 | Page 9, 10 |
| Table S10 | Page 11 |

**Table S2.** ANOVA results for the interaction between race and country.

| **Variation source** | **Sum of Squares** | **Mean Square** | **DF (num)** | **DF (den)** | **F** | **p** |
| --- | --- | --- | --- | --- | --- | --- |
| **Race** | 0.08 | 0.027 | 3 | 7.65 | 1.03 | 0.430 |
| **Country** | 0.11 | 0.115 | 1 | 7.80 | 4.34 | 0.072 |
| **Race x Country** | 0.03 | 0.009 | 3 | 7.65 | 0.33 | 0.807 |

Abbreviations: DF (num) = Numerator Degrees of Freedom, DF (den) = Denominator Degrees of Freedom.

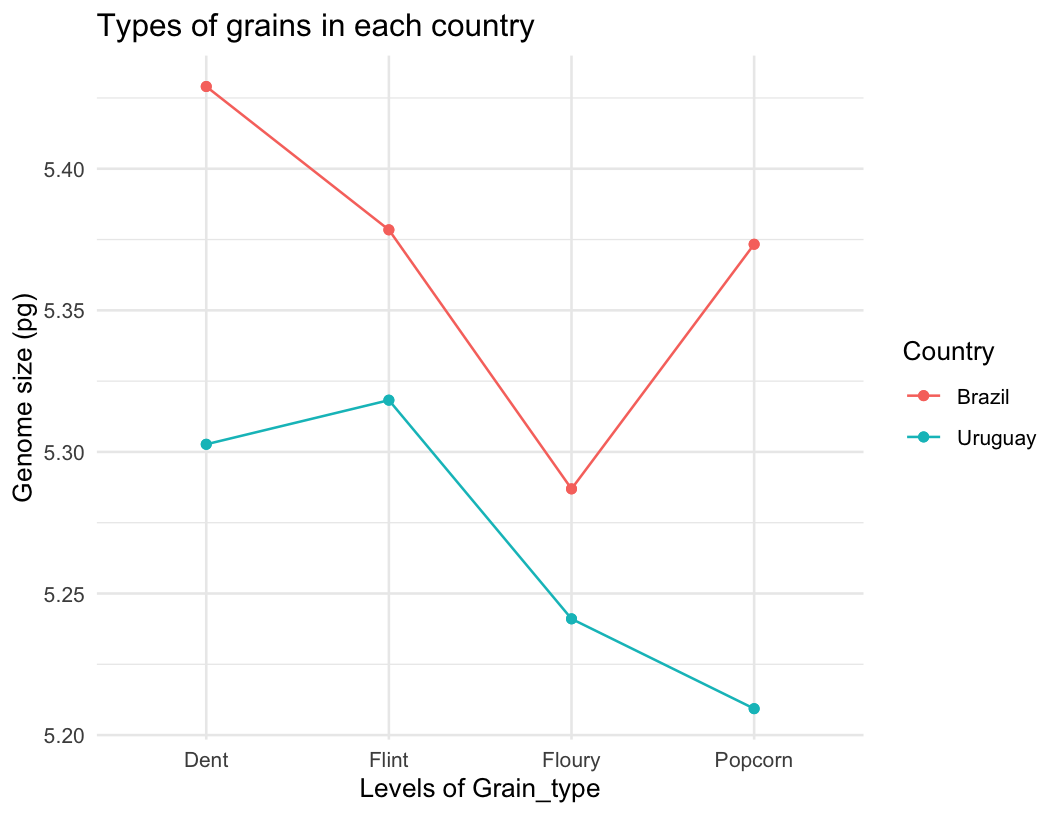

**Figure S1.** Genome size (2C values) comparisons between Uruguay and Brazil across maize grain type. Each colored line represents a different maize grain type.

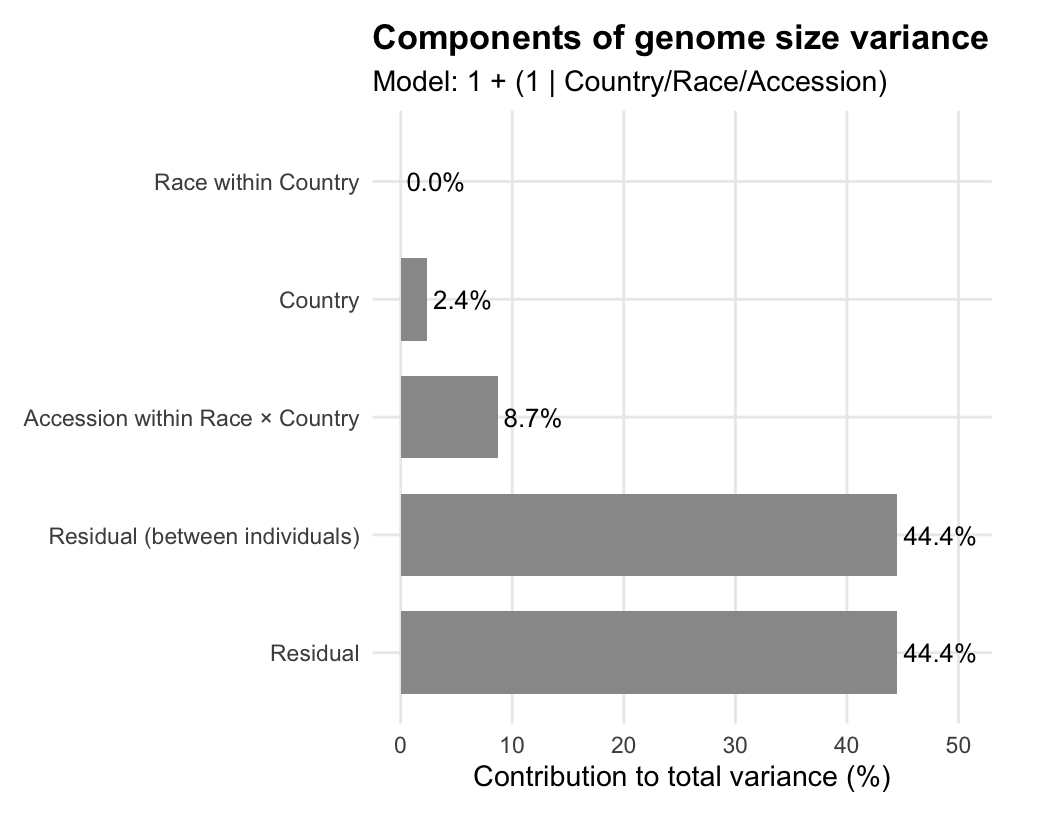

**Figure S2.** Components of genome size variance.

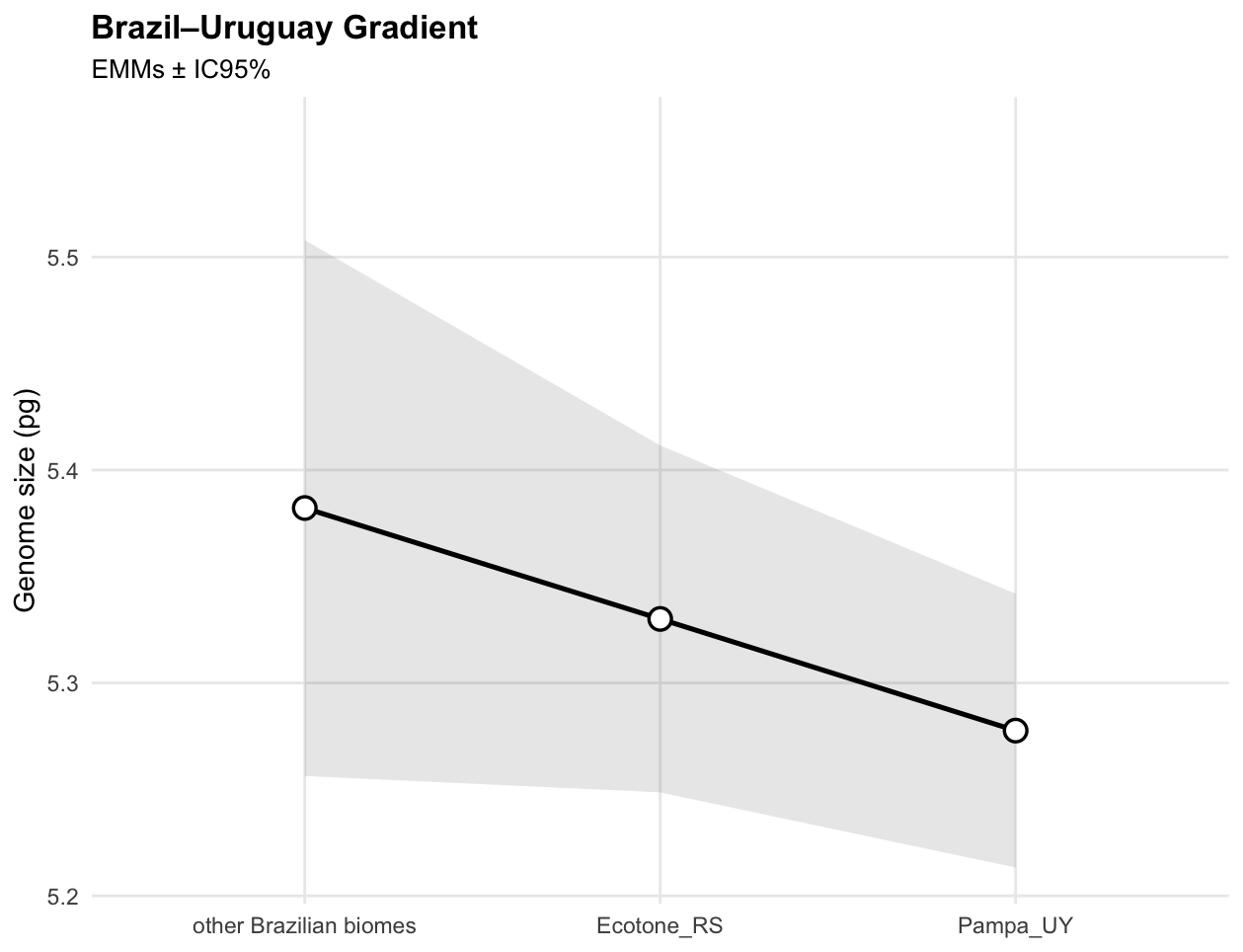

**Figure S3.** Ecological gradient of genome size between Brazil and Uruguay (F_2, 12.04_ = 1.55, p = 0.250). Each point on the graph shows the estimated marginal mean of the genome size within each region, and the gray shading indicates the confidence interval (95%).

**Table S3.** Pairwise contrasts of the Blanco Dentado race.

| **Contrast** | **Ratio** | **SE** | ***z*** | ***p*-value** |
| --- | --- | --- | --- | --- |
| **RSG4A / UYEB1B** | 1.37 | 0.150 | 2.906 | 0.010* |
| **RSG4A / UYER3A** | 1.57 | 0.178 | 4.006 | < 0.001* |
| **UYEB1B / UYER3A** | 1.15 | 0.131 | 1.194 | 0.457 |

P value adjustment: tukey method for comparing a family of 3 estimates. Tests are performed on the log scale. * indicate statistically significant differences (p < 0.05). Abbreviations: SE = Standard Error.

**Table S4.** Fixed effects from the generalized linear mixed model that predicts the pattern of variation in the number of knobs, adjusted for Brazil.

| **Term/Predictor** | **Estimate (β)** | **SE** | **Statistic** | ***p*-value** |
| --- | --- | --- | --- | --- |
| (Intercept) | 3.34 | 0.08 | 43.60 | < 0.001 |
| Complejo Cateto | -0.11 | 0.09 | -1.17 | 0.241 |
| Complejo Moroti-Caingang | -0.26 | 0.09 | -2.99 | 0.003 |
| Pisingallo | -0.21 | 0.10 | -2.06 | 0.039 |

Abbreviations: SE = Standard Error.

**Table S5.** Fixed effects from the generalized linear mixed model that predicts the pattern of variation in the number of knobs, adjusted for Uruguay.

| **Term/Predictor** | **Estimate (β)** | **SE** | **Statistic** | ***p*-value** |
| --- | --- | --- | --- | --- |
| (Intercept) | 2.96 | 0.06 | 52.05 | <0.001 |
| Complejo Cateto | 0.16 | 0.07 | 2.09 | 0.036 |
| Complejo Moroti-Caingang | 0.19 | 0.10 | 1.86 | 0.063 |
| Pisingallo | 0.22 | 0.08 | 2.88 | 0.004 |

Abbreviations: SE = Standard Error.

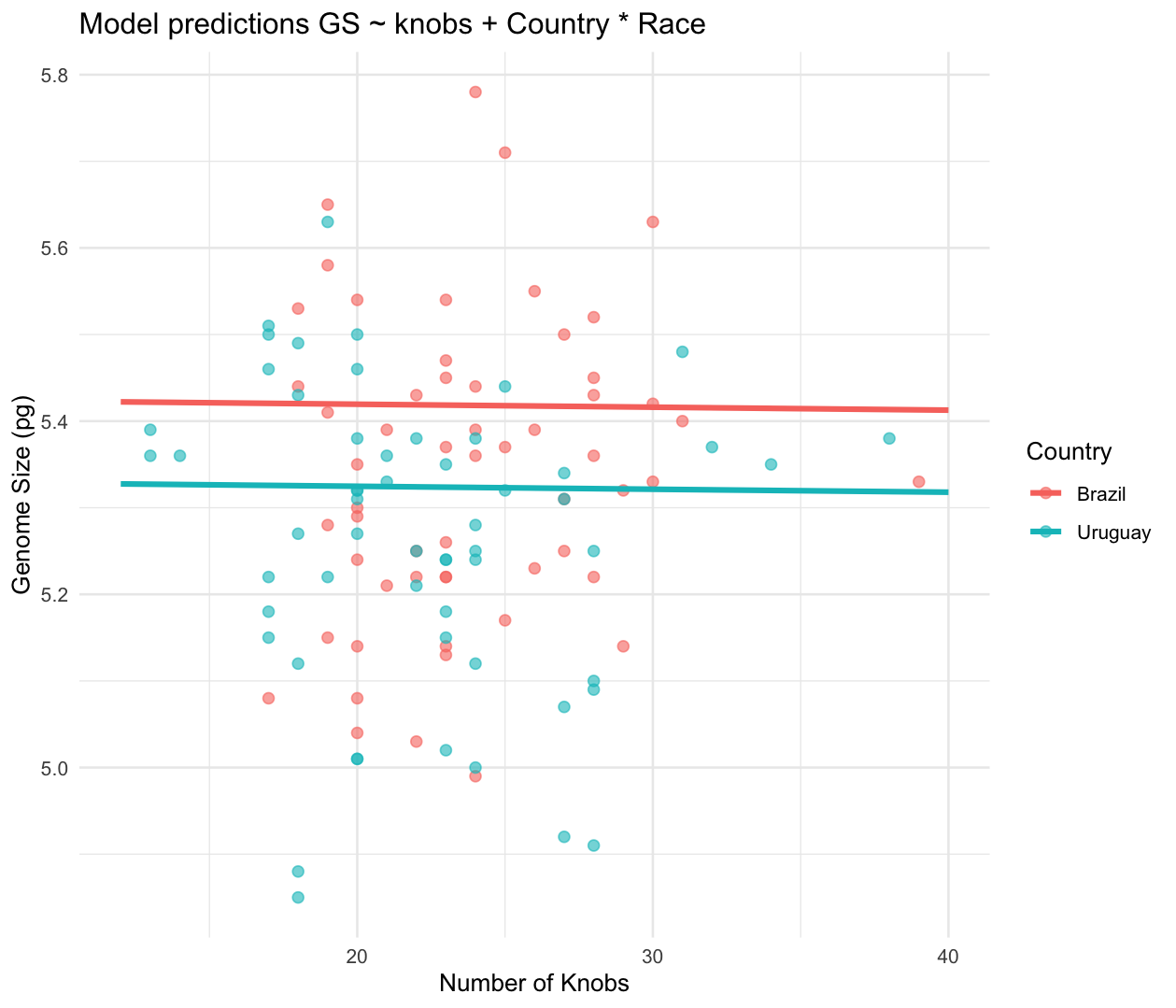

**Figure S4.** Linear relationship between the knobs numbers and genome size.

**Table S6.** Summary of linear and linear mixed-effects models evaluating the relationship between knob numbers and genome size under different covariate controls.

| **Model ID** | **Model Specification** | **Estimate (β)** | **SE** | **Statistic** | ***P*-value** | **AIC** | **R²** |
| --- | --- | --- | --- | --- | --- | --- | --- |
| M0 | Genome ~ Knobs | 0.0018 | 0.004 | 0.49 | 0.624 | -65.9 | 0.002 |
| M2 | Genome ~ Race + Knobs | 0.0013 | 0.004 | 0.37 | 0.713 | -68.4 | 0.074 |
| M4 | Genome ~ Country + Knobs | 0.0003 | 0.004 | 0.07 | 0.943 | -69.5 | 0.050 |
| M5 | Genome ~ Knobs + Country * Race | -0.0008 | 0.004 | -0.21 | 0.832 | -73.8 | 0.177 |
| M7 (LMM) | Genome ~ Knobs + Country * Race + (1\|Race : Accession) | -0.0003 | 0.004 | -0.09 | 0.929 | -30.7 | 0.275 |

**Table S7.** Fixed effect from the generalized linear mixed model (GLMM) predicting the relationship between knob numbers and latitude.

| **Fixed Effect** | **Estimate (β)** | **SE** | ***z*** | ***p*-value** |
| --- | --- | --- | --- | --- |
| (Intercept) | 3.13 | 0.03 | 117.04 | <0.001 |
| Latitude | 0.04 | 0.03 | 1.62 | 0.104 |

**Table S8.** Fixed effect from the generalized linear mixed model (GLMM) predicting the relationship between knob numbers, latitude and genome size.

| **Fixed Effect** | **Estimate (β)** | **SE** | ***z*** | ***p*-value** |
| --- | --- | --- | --- | --- |
| (Intercept) | 3.23 | 0.65 | 4.96 | <0.001 |
| Latitude | 0.04 | 0.03 | 1.62 | 0.106 |
| Genome size | -0.02 | 0.12 | -0.15 | 0.877 |

**Table S9.** Genomic proportions (in Mb) of each repeat class and families of maize landraces from Brazil and Uruguay annotated by Repeat Explorer.

PENSAR SOBRE LOS NOMBRES DE LAS ACCESIONES

| **Repeat Classification** | **PBF2A1** | **RSG4A11** | **RSG412** | **UYNN4D11** | **UYNN4D12** |
| --- | --- | --- | --- | --- | --- |
| All/repeat/mobile_element/Class_I/LINE | 1.44 | 1.08 | 1.32 | 1.17 | 1.10 |
| All/repeat/mobile_element/Class_I/LTR | 22.97 | 22.00 | 23.09 | 21.22 | 22.82 |
| All/repeat/mobile_element/Class_I/LTR/Ty1_copia/Ale | 3.50 | 2.79 | 3.21 | 3.13 | 2.80 |
| All/repeat/mobile_element/Class_I/LTR/Ty1_copia/Angela | 6.09 | 5.79 | 6.19 | 5.67 | 6.22 |
| All/repeat/mobile_element/Class_I/LTR/Ty1_copia/Bianca | 1.38 | 2.13 | 1.16 | 1.27 | 1.30 |
| All/repeat/mobile_element/Class_I/LTR/Ty1_copia/Ivana | 1.82 | 1.53 | 2.36 | 1.63 | 1.85 |
| All/repeat/mobile_element/Class_I/LTR/Ty1_copia/SIRE | 513.28 | 520.56 | 527.45 | 518.50 | 519.27 |
| All/repeat/mobile_element/Class_I/LTR/Ty1_copia/TAR | 12.36 | 11.43 | 12.52 | 11.91 | 11.79 |
| All/repeat/mobile_element/Class_I/LTR/Ty1_copia/Tork | 0.43 | 0.29 | 0.55 | 0.26 | 0.40 |
| All/repeat/mobile_element/Class_I/LTR/Ty3_gypsy/chromovirus/CRM | 28.80 | 25.84 | 27.45 | 28.53 | 28.39 |
| All/repeat/mobile_element/Class_I/LTR/Ty3_gypsy/chromovirus/Tekay | 476.74 | 479.73 | 477.30 | 469.86 | 467.30 |
| All/repeat/mobile_element/Class_I/LTR/Ty3_gypsy/non-chromovirus/OTA/Tat/Ogre | 18.99 | 18.39 | 18.47 | 18.16 | 19.53 |
| All/repeat/mobile_element/Class_I/LTR/Ty3_gypsy/non-chromovirus/OTA/Tat/Retand | 505.33 | 512.64 | 520.58 | 530.13 | 515.54 |
| All/repeat/mobile_element/Class_II/Subclass_1/TIR/EnSpm_CACTA | 35.85 | 33.61 | 37.96 | 33.41 | 32.39 |
| All/repeat/mobile_element/Class_II/Subclass_1/TIR/MuDR_Mutator | 1.04 | 1.31 | 1.37 | 1.40 | 1.02 |
| All/repeat/mobile_element/Class_II/Subclass_1/TIR/PIF_Harbinger | 0.14 | 0.32 | 0.30 | 0.44 | 0.34 |
| All/repeat/mobile_element/Class_II/Subclass_2/Helitron | 1.05 | 1.43 | 1.34 | 1.21 | 1.35 |
| All/repeat/rDNA/45S_rDNA | 43.42 | 41.29 | 21.40 | 45.00 | 32.63 |
| All/repeat/rDNA/5S_rDNA | 1.04 | 0.96 | 0.59 | 0.67 | 1.14 |
| All/repeat/satellite/156/CentC | 6.01 | 6.91 | 7.15 | 9.78 | 7.65 |
| All/repeat/satellite/180/knob180 | 147.98 | 131.17 | 110.35 | 113.64 | 108.24 |
| All/repeat/satellite/358/knobTR1 | 29.42 | 25.90 | 13.17 | 15.06 | 11.14 |
| All/repeat/satellite/385/cent4 | 0.21 | 0.39 | 0.47 | 0.21 | 0.18 |
| All/repeat/satellite/531/telomeric sequence | 1.75 | 1.76 | 1.70 | 1.72 | 1.72 |
| unclassified | 61.62 | 61.11 | 62.26 | 61.74 | 60.81 |
| **Total repetitive DNA** | 1,922.64 | 1,910.34 | 1,879.73 | 1,895.72 | 1,856.92 |

**Table S10.** Races and landraces exhibiting variation in the number of B chromosomes.

| **Country** | **Race** | **Accession** | **B Chromosomes** |
| --- | --- | --- | --- |
| **Uruguay** | Pisingallo | URZM03013 | 2 |
| **Uruguay** | Pisingallo | URZM03013 | 2 |
| **Uruguay** | Pisingallo | URZM03013 | 1 |
| **Uruguay** | Pisingallo | URZM03013 | 2 |
| **Uruguay** | Pisingallo | URZM1111 | 1 |
| **Uruguay** | Pisingallo | URZM1111 | 1 |
| **Uruguay** | Pisingallo | URZM1111 | 1 |
| **Uruguay** | Cateto_Sulino | UYEN1A | 1 |
| **Uruguay** | Cateto_Sulino | UYEN1A | 1 |
| **Uruguay** | Cateto_Sulino | UYEN1A | 1 |
| **Uruguay** | Cateto_Sulino | UYEN1A | 1 |
